# A parsimonious, age-invariant, and psychometrically validated neural signature of task-control network functioning

**DOI:** 10.64898/2026.10.08.757721

**Authors:** K.J. Paige, M.F. Molloy, M. Angstadt, L.W. Hyde, C. Sripada, M.M. Heitzeg, A.S. Weigard

**Affiliations:** University of Michigan, Psychiatry, Ann Arbor, Michigan; The Pennsylvania State University, Psychology, University Park, Pennsylvania; University of Michigan, Psychology, Ann Arbor, Michigan

## Abstract

Neuroimaging has vast potential for identifying brain features that support individual and developmental differences in cognition, but challenges related to the dimensionality, unreliability, and unknown developmental properties of neuroimaging measures prevent this from being realized. We overcome these challenges by applying latent variable modeling to demand-related responses in the “task-control” frontoparietal and dorsal attention networks (FPN, DAN) within the Adolescent Brain Cognitive Development Study and Human Connectome Project. Task-control activations are well-captured by a bifactor structure with a general factor spanning FPN/DAN and network-specific factors that is remarkably invariant across middle childhood to early adolescence and largely generalizes to adulthood. Trait-state-occasion modeling reveals a general task-control trait that shows robust associations with cognitive abilities, clinical symptoms, and grades. Repeated neuroimaging substantially increases associations. Findings establish a parsimonious neural signature that generalizes across samples, is stable across development, and has broad utility for understanding developmental and individual differences in goal-directed behavior.

**Graphical Abstract:** 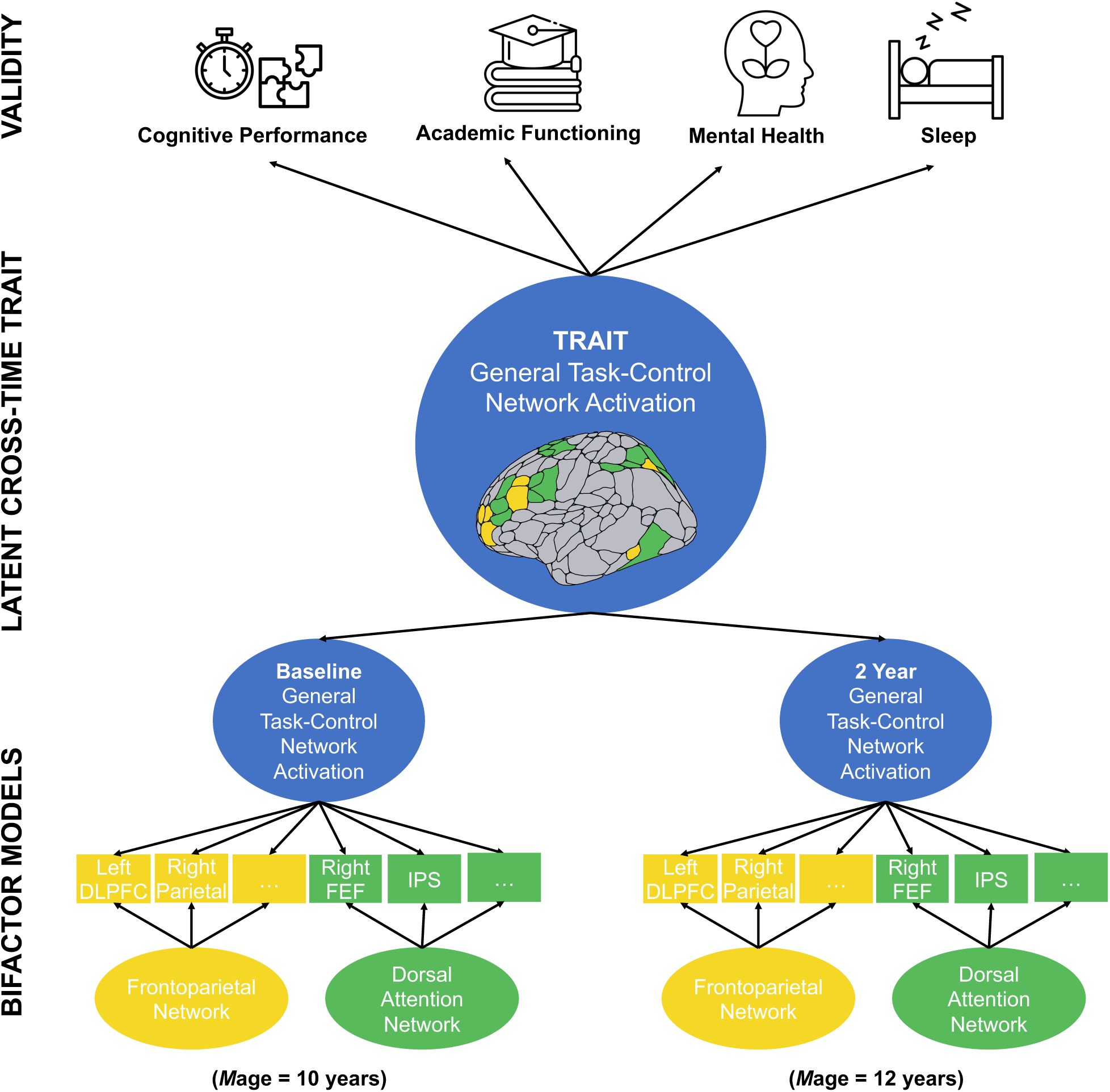

## Introduction

Individual differences in cognitive abilities, including processing speed, attention, problem solving, planning, and inhibitory control, are broadly associated with better mental health, educational achievement, and life success^1–3^. Longitudinal research demonstrates that difficulties in cognitive abilities in middle childhood predict a sequela of long-term life outcomes, including mental and physical health, substance use problems, personal finances, and legal problems^3–7^. Given the well-documented importance of healthy cognitive development in reducing risk for negative outcomes and supporting adaptive functioning across the lifespan, understanding the features of the brain that underlie individual and developmental differences in cognitive functioning is critical for public health and has been a major goal of contemporary neuroscience^8–13^.

In humans, neuroimaging methods such as functional magnetitic resonance imaging (fMRI) have been the dominant measurement modality used to characterize the neural underpinnings of individual and developmental differences in cognition. These methods show immense promise for identifying neural features that underlie the development of individuals’ cognitive functioning and goal-directed behavior^14,15^. Further, if these features can be measured reliably and are related to real-word outcomes (such as psychopathology and academic functioning), they could significantly advance translational applications. A robust and meaningful individual-level neural signature could better characterize risk for psychopathology and inform efforts to facilitate adaptive neurocognitive development in youth.

Yet, despite the promise of this approach, the use of fMRI to identify reliable individual differences has encountered major challenges^9,16–19^. Specifically, measures of the functioning of individual brain regions typically have poor temporal stability and weak associations with relevant behavioral measures^16–18^. Recent work that has sought to improve the reliability and generalizability of fMRI measures relevant to cognitive ability using data-driven, predictive models applied to high-dimensional neuroimaging data, typically from resting-state fMRI^20–23^. These approaches have been instrumental in demonstrating that variance related to cognitive ability is widely distributed across the brain and particularly implicates “task-control” frontoparietal (FPN) and dorsal attention (DAN) networks^22,24–28^ (**Figure 1**). However, several critical limitations remain.

**Figure 1.**
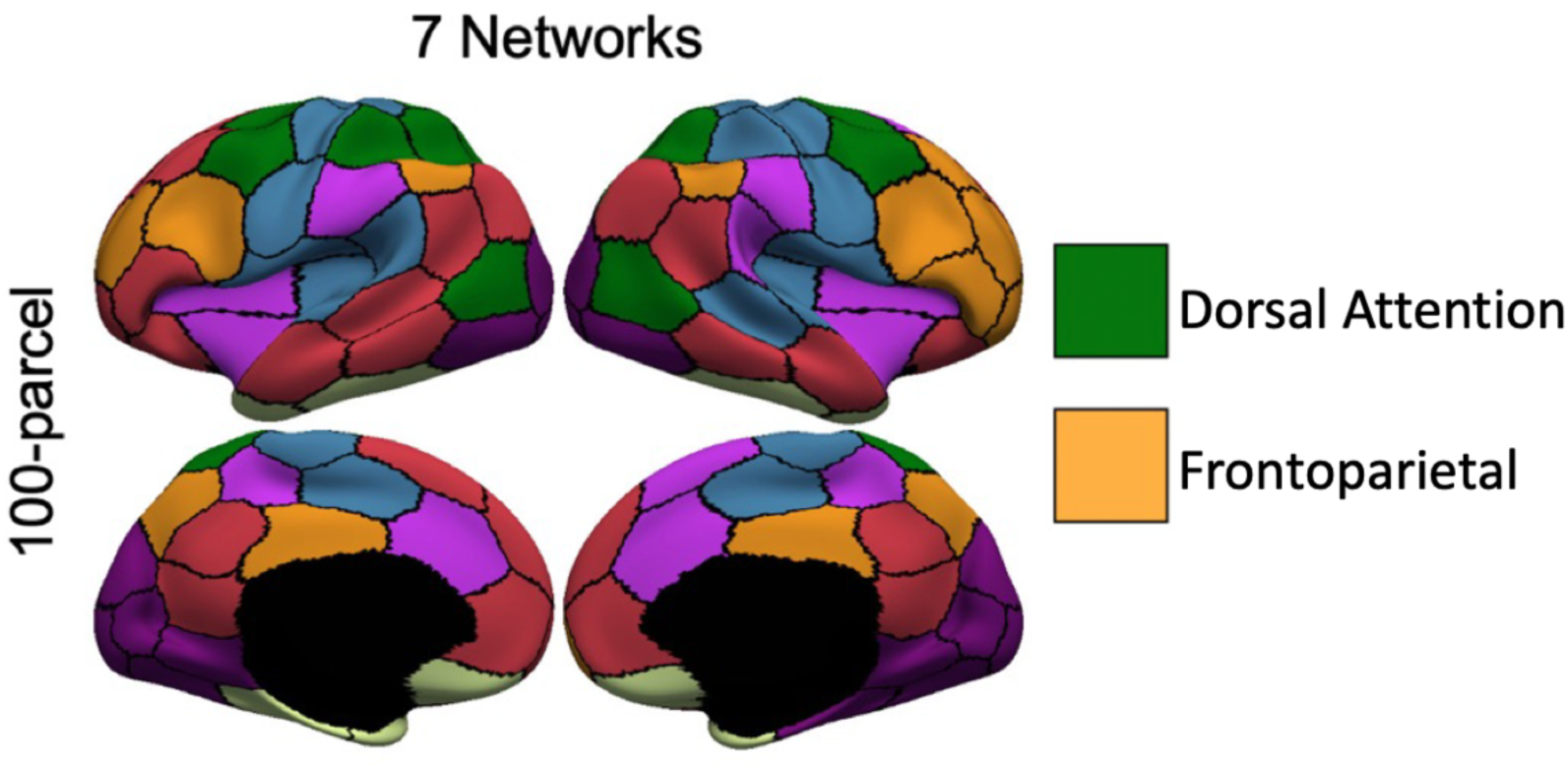
*Schaefer 100-parcel, 7-network cortical parcellation*. Cortical regions are parcellated into 100 regions of interest spanning seven canonical functional networks. Dorsal Attention and Frontoparietal Networks are depicted in green and yellow, respectively.

First, from a practical standpoint, these high-dimensional predictive modeling approaches are cumbersome and resource-intensive: they require very large sample sizes^21^, long scanning sessions to optimize reliability (i.e., approximately one hour)^22^, and multivariate models with thousands of parameters (i.e., beta weights estimated for each neural feature) that have a strong tendency overfit data in specific samples and fail to generalize to others^29,30^. These practical considerations substantially limit the scalability and reproducibility of neural metrics across labs, samples, and populations. Second, the parametric complexity and largely atheoretical nature of data-driven approaches significantly constrains their interpretability. Because such models are optimized for predictive accuracy rather than explanation, it can be difficult to clearly map their findings of diffuse, high-dimensional patterns of neural data onto parsimonious explanatory theories of the specific neural mechanisms support cognition and behavior^31–33^. Third, these data-driven approaches do little to clarify the reliability, developmental stability, or trait/state properties of the diffuse neural features they identify. This is a major limitation because understanding developmental change requires that neural features are measured consistently and accurately across ages. Taken together, these limitations highlight the need for scalable, interpretable, and psychometrically informed approaches to identifying neural signatures of cognitive ability^16,34^. The current study addresses these gaps by leveraging well-established psychometric methods to clarify key measurement properties of a parsimonious and interpretable signature of brain network dynamics that is strongly predictive of individuals’ cognitive ability.

Building on recent work demonstrating that responses to cognitive demand in the FPN and DAN are strongly linked to cognitive performance^35,36^, we apply psychometric methods across two large fMRI datasets spanning middle childhood through young adulthood to develop a parsimonious and stable measure of individuals’ task-control network functioning (**Figure 1**). We establish task-control network responses to cognitive demand as a unitary construct supporting cognitive development and quantify its measurement accuracy, stability across development, and trait- and occasion-level associations with major behavioral outcomes. For the first time, our study reveals a parsimonious, reproducible, and psychometrically robust neural signature of task-control network functioning that has broad applications for characterizing the neural mechanisms that drive developmental and individual differences in goal-directed cognition and modeling their maturational trajectories.

## Results

### Responses to cognitive demand in FPN and DAN regions predominantly reflect a single general “task-control” factor

Using data from the baseline (*M*age=10.01 years; *n*=4862) and 2-year follow-up (*M*age=11.99; *n*=3648) sample from Adolescent Brain Cognitive Development^SM^ Study (ABCD Study®)^37^ and the baseline sample (*M*age=28.78; *n*=1080) from the Human Connectome Project (HCP)^38^, we sought to assess the suitability of a factor structure for FPN and DAN activations that reflected two key assumptions grounded in prior work; 1) the large body of evidence from resting state and task-based neuroimaging suggesting that the FPN and DAN represent independent networks with dissociable functions^39–42^, and 2) evidence from a prior study^36^ indicating that the FPN and DAN show similar responses to n-back load and similar relations to cognition in this context, suggesting a general component that explains variance across both task-control networks. Such a structure is well-described by a bifactor model^43^ assuming that variance in each FPN and DAN region is jointly explained by a network-specific (FPN or DAN) latent variable as well as by an orthogonal general latent variable that accounts for common variance across both networks (**Figure 2**).

**Figure 2.**
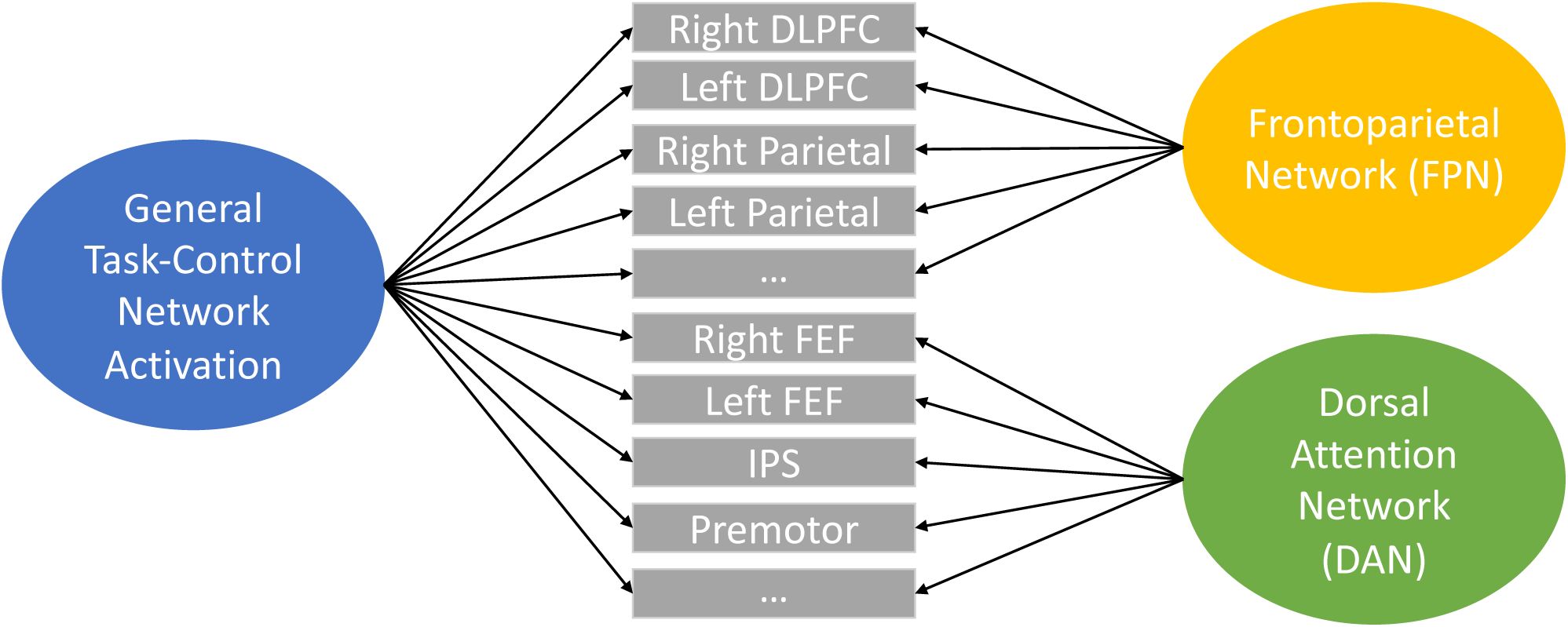
*Conceptual Figure of a Bifactor Model of General Task-Control Network Activation*. Bifactor modeling allows for separation of variance general to, or shared by, neural activation across the task-control networks from variance unique to the frontoparietal and dorsal attention networks. DLPFC=dorsolateral prefrontal cortex. FED=frontal eye fields. IPS=intraparietal sulcus.

Bifactor models whereby all ROIs loaded onto a general factor of task-control neural activation alongside two network-specific factors for the FPN and DAN converged and showed acceptable fit across all neuroimaging assessments (**Supplementary Table S1**).

A dominant and reliable general factor (ω range=0.94 to 0.96; H range=0.93 to 0.96) accounted for substantial individual differences in task-control network activation and had strong factor loadings across regions spanning the FPN and DAN, across all three assessments (**Table 1**). The general factors of task-control network activation accounted for the majority of explainable variance across development (Explained common variance range=0.66 to 0.73; ωH range=0.77 to 0.84), indicating that activation across these distributed regions was largely captured by a common underlying dimension. Together, these findings support a robust, general neural dimension of task-control activation spanning the FPN and DAN.

**Table 1.**
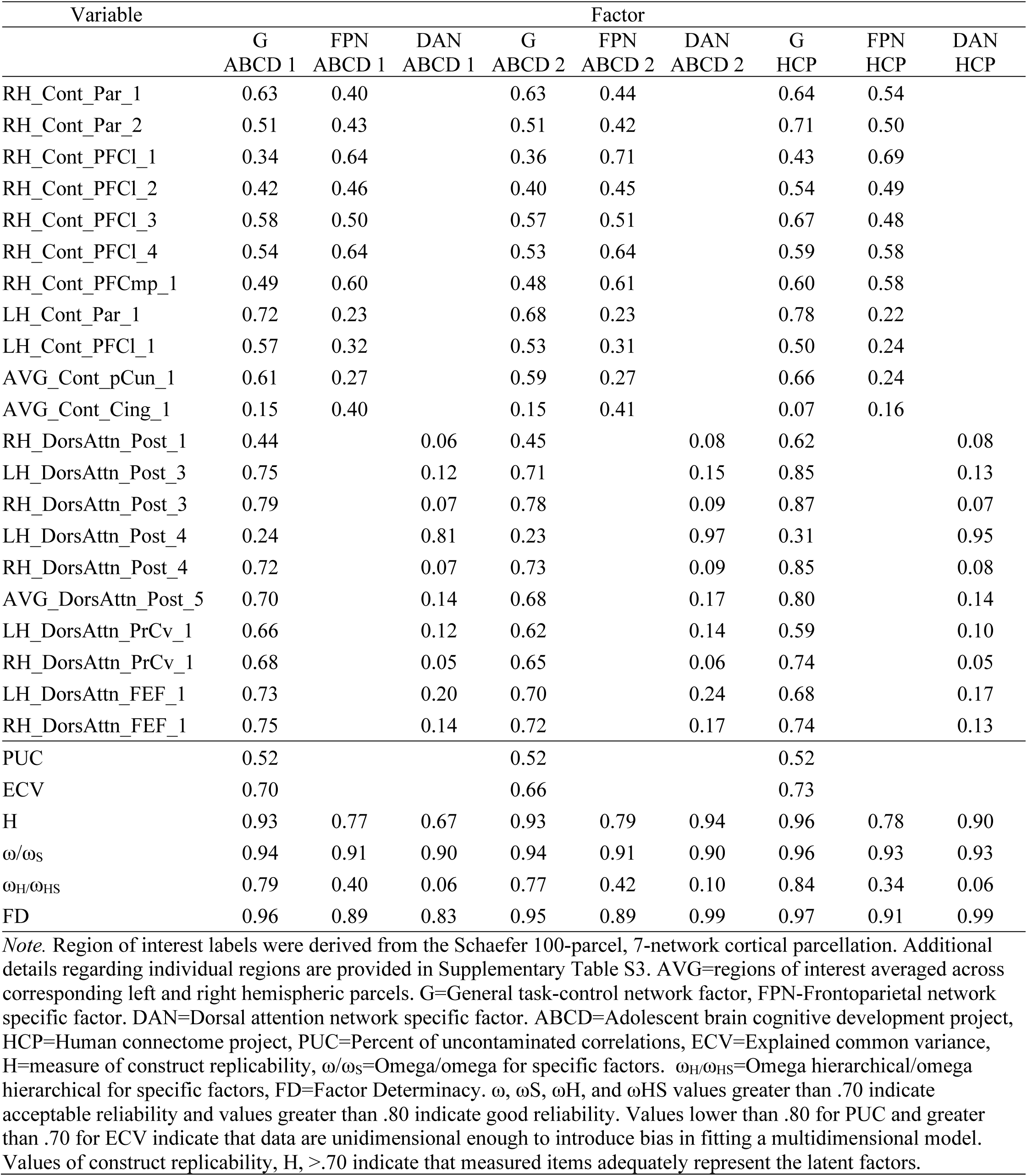
Standardized factor loadings for bifactor models imposing partial metric invariance across ABCD and HCP.

Although the general factor accounted for substantial variance across regions, bifactor alternative fit statistics indicated that activation across the task-control networks during cognitive demand also reflected meaningful network-specific variation (Percent of Uncontaminated Correlation values=0.52 across assessments; **Table 1**). This pattern supports our use of bifactor models, which distinguish between what these networks have in common and what is unique to each network. With respect to the specific factors, the FPN showed evidence of capturing meaningful information beyond the general task-control factor (H range=0.77 to 0.79; ω_HS_ range=0.34 to 0.42). The DAN specific factors were also reasonably well-defined (H range=0.67 to 0.94), but they accounted for very little reliable variance (ω_HS_ range=0.06 to 0.10). This pattern was also evident in the factor loadings: the specific FPN factors were characterized by generally strong loadings across time whereas the specific DAN factors mostly demonstrated weak loadings. Taken together, these results suggest that the FPN contributed meaningful network-specific information beyond what was shared across regions, whereas most of the reliable variance in the DAN was captured by the general factor.

Next, we combined these measurement models across ages. The confirmatory factor model with three correlated task-control bifactor models (one bifactor model at each assessment) provided good fit (**Table 2**; Configural), supporting configural measurement invariance. In summary, results supported a consistent bifactor structure of task-control neural activation, in which neural activation across the FPN and DAN network load onto a general factor and network-specific factors across middle childhood, early adolescence, and young adulthood. These data exhibited a dominant and reliable general factor that accounted for substantial individual differences in task-control network activation across development.

**Table 2.**
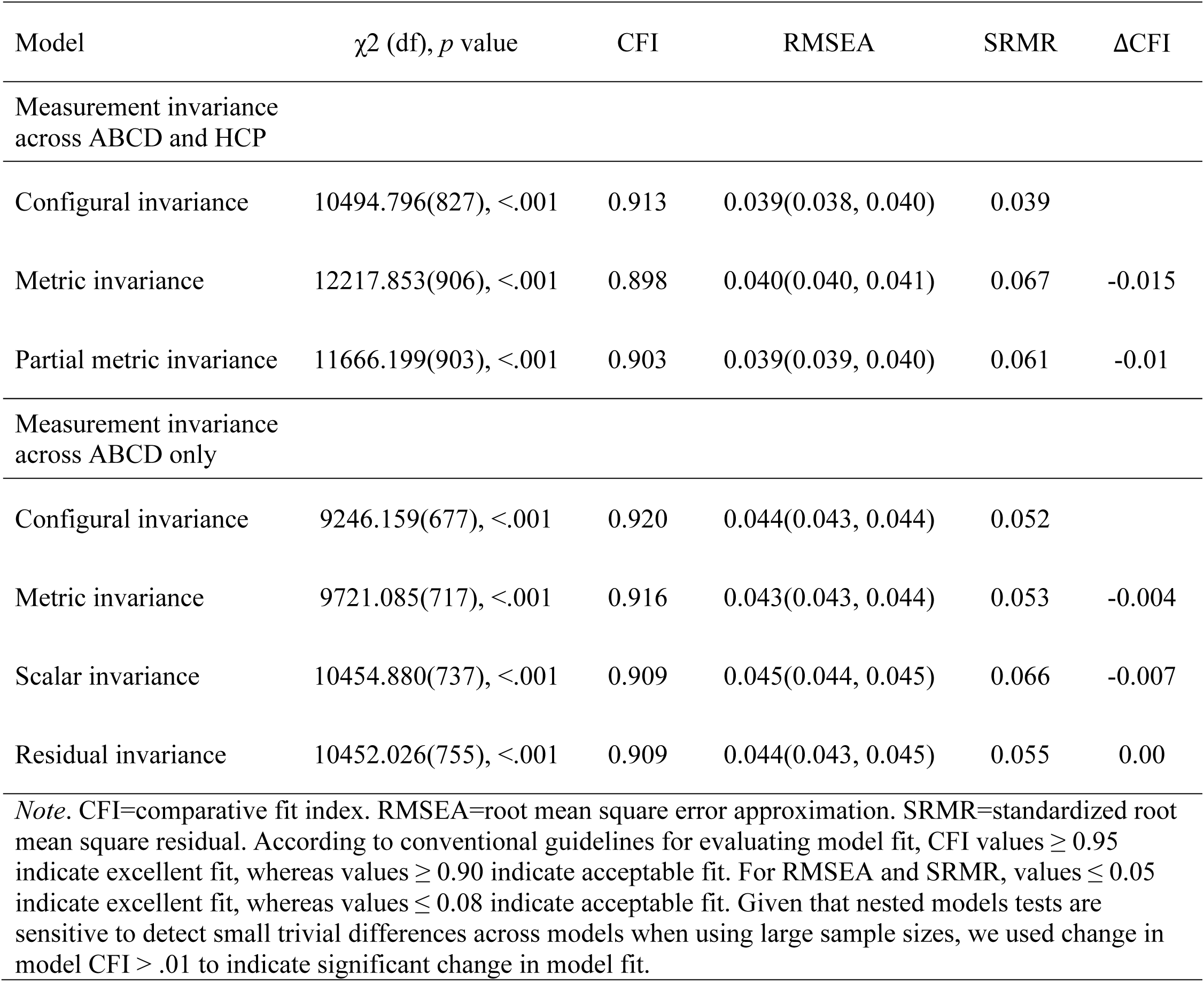
Fit statistics for bifactor models testing measurement invariance.

| Model | $\chi^2$ (df), $p$ value | CFI | RMSEA | SRMR | ΔCFI |
| --- | --- | --- | --- | --- | --- |
| Measurement invariance across ABCD and HCP |  |  |  |  |  |
| Configural invariance | 10494.796(827), <.001 | 0.913 | 0.039(0.038, 0.040) | 0.039 |  |
| Metric invariance | 12217.853(906), <.001 | 0.898 | 0.040(0.040, 0.041) | 0.067 | -0.015 |
| Partial metric invariance | 11666.199(903), <.001 | 0.903 | 0.039(0.039, 0.040) | 0.061 | -0.01 |
| Measurement invariance across ABCD only |  |  |  |  |  |
| Configural invariance | 9246.159(677), <.001 | 0.920 | 0.044(0.043, 0.044) | 0.052 |  |
| Metric invariance | 9721.085(717), <.001 | 0.916 | 0.043(0.043, 0.044) | 0.053 | -0.004 |
| Scalar invariance | 10454.880(737), <.001 | 0.909 | 0.045(0.044, 0.045) | 0.066 | -0.007 |
| Residual invariance | 10452.026(755), <.001 | 0.909 | 0.044(0.043, 0.045) | 0.055 | 0.00 |
| *Note.* CFI=comparative fit index. RMSEA=root mean square error approximation. SRMR=standardized root mean square residual. According to conventional guidelines for evaluating model fit, CFI values $\geq 0.95$ indicate excellent fit, whereas values $\geq 0.90$ indicate acceptable fit. For RMSEA and SRMR, values $\leq 0.05$ indicate excellent fit, whereas values $\leq 0.08$ indicate acceptable fit. Given that nested models tests are sensitive to detect small trivial differences across models when using large sample sizes, we used change in model CFI $> .01$ to indicate significant change in model fit. | | | | | |

### Measurement invariance indicates that developmental change in task-control network activations is largely quantitative rather than qualitative

Next, we proceeded to test stricter forms of measurement invariance to examine the consistency of the factor structure across ages, independent samples, and task designs. We first explored measurement invariance testing of bifactor models in the ABCD subsample across baseline and 2-year follow-up assessments given the fact that invariance would not necessarily be expected between ABCD and HCP due to differences in sample characteristics, task design, and imaging parameters. Metric invariance (equal factor loadings) tests whether the strength of the relationship between each observed ROI and the latent variable is consistent across groups or time points, allowing comparisons of associations involving the latent variable. Imposing equal factor loadings across these two repeated measures in ABCD did not lead to a significant decrement in model fit (**Table 2**), supporting full metric invariance. Next, we examined scalar invariance by imposing equal intercepts. Scalar invariance indicates that if two individuals from different developmental periods (e.g., a 10-year-old and 12-year-old) have the same level of the latent variable, they will, on average, have the same observed scores on the ROIs that contribute to this latent variable. Scalar invariance is essential to infer that differences in scores across groups and/or developmental periods reflect true differences in the underlying construct (e.g., task-control network activation). Results supported scalar invariance. Finally, strict invariance (equal residual variances) tests whether the amount of measurement error is consistent across groups or time points, providing the strongest evidence that the construct is being measured in the same way. Remarkably, strict measurement invariance across the two ABCD assessments was also supported. The final model provided an acceptable fit to the data (**Table 1**). This most stringent level of measurement invariance allows us to confidently interpret longitudinal change in these metrics. In other words, changes in scores across this two year measurement interval reflect quantitative developmental change rather than differences in measurement properties over time.

We then tested metric invariance across both ABCD waves and HCP, examining whether each ROI contributes to the latent general and network-specific factors to the same degree in independent samples across distinct developmental periods. Partial metric invariance was supported after freeing equality constraints on two loadings on the general factor (right inferior parietal lobule, right medial posterior prefrontal cortex) and one loading on the specific FPN factor (right medial posterior prefrontal cortex) in HCP, as these regions had stronger loadings in the HCP (**Table 1**). Thus, results indicated that the vast majority of ROIs contribute to the latent constructs (the general factor as well as specific DAN and FPN factors) with the same strength across middle childhood, early adolescence, and young adulthood.

### The general factor represents a stable and valid signature of task-control network functioning across developmental periods spanning middle childhood to early adulthood

To examine stability, we regressed the general and specific latent variables from ABCD 2-year follow-up onto the general and specific latent variables from ABCD baseline. In the configural model, the general task-control network activation factor demonstrated stability (β=0.29) across the two-year interval; the specific FPN (β=0.23) and DAN (β=0.10) factors demonstrated lower stability. Imposing strict measurement invariance strengthened the stability of the general task-control network activation factor (β=0.35) as well as the specific FPN (β=0.26) and DAN (β=0.27) factors. These stability coefficients are substantial given the relatively long measurement interval of 2 years and exceed those previously reported for individual ROIs in this task and sample^17^.

To test validity, we next examined within-time associations between the general task-control network factors, specific FPN and DAN factors, and a wide array of criterion measures. Informed by a meta-analysis of effect sizes in individual-differences research, correlations of 0.10, 0.20, and 0.30 can be interpreted as small, medium, and large, respectively^44^. Large, positive associations between our neural measures and performance on cognitive tasks would provide strong evidence of convergent validity. Following prior work^36,45–47^, we estimated the diffusion model’s drift rate parameter from n-back data to provide a reliable and valid measure of individuals’ general efficiency of cognitive performance on the task performed concurrently with fMRI data collection. We also evaluated criterion relations with drift rate from a non-concurrent task (stop-signal task; SST) and several more traditional cognitive measures administered outside of the scanner. All models included in-scanner head motion, and baseline age, sex, and an indicator of socioeconomic status (child opportunity index, “coi_z”^48^ for ABCD; education for HCP). Model fit statistics for all validity test models are reported in **Supplementary Table S2**. In HCP (**Figure 3**), the general factor of task-control activation demonstrated a large positive association with concurrent drift rate (β=0.42) and small-to-medium positive associations with other cognitive measures (β range=0.07 to 0.23). The specific FPN factor demonstrated a small positive association with concurrent drift rate (β=0.15) only. The specific DAN factor was negatively associated with cognitive abilities, and all effects were small in magnitude.

**Figure 3.**
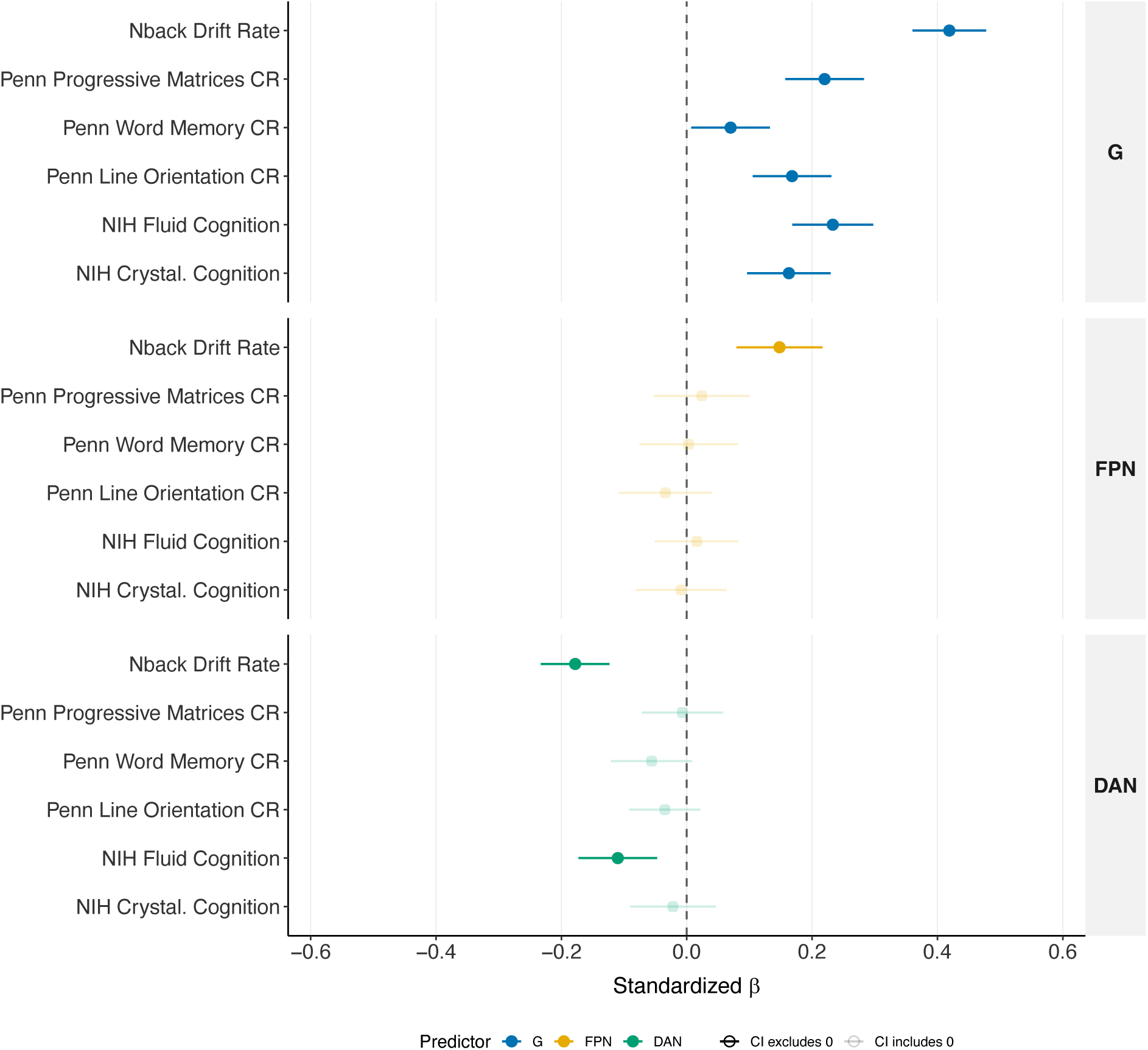
*Forest plot depicting criterion validity associations between general task-control network activation, network specific activation, and cognitive measures in the Human Connectome Project sample.* G=general task-control network activation latent factor. FPN=specific frontoparietal network latent factor. DAN=specific dorsal attention network latent factor. CR=Correct response. NIH=NIH toolbox cognition battery measure. Crystal.=Crystallized.

Regarding ABCD (**Figure 4**), the general factor showed large positive associations with concurrent n-back drift rate (β_T1_=0.45; β_T2_=0.35) and small-to-medium positive associations with other cognitive tasks. Consistent with HCP, the specific FPN factor demonstrated a small positive association with concurrent drift rate at 2-year follow-up (β=0.09). Other relations with the FPN and DAN factors were not significant or small and in the opposite of the expected direction.

**Figure 4.**
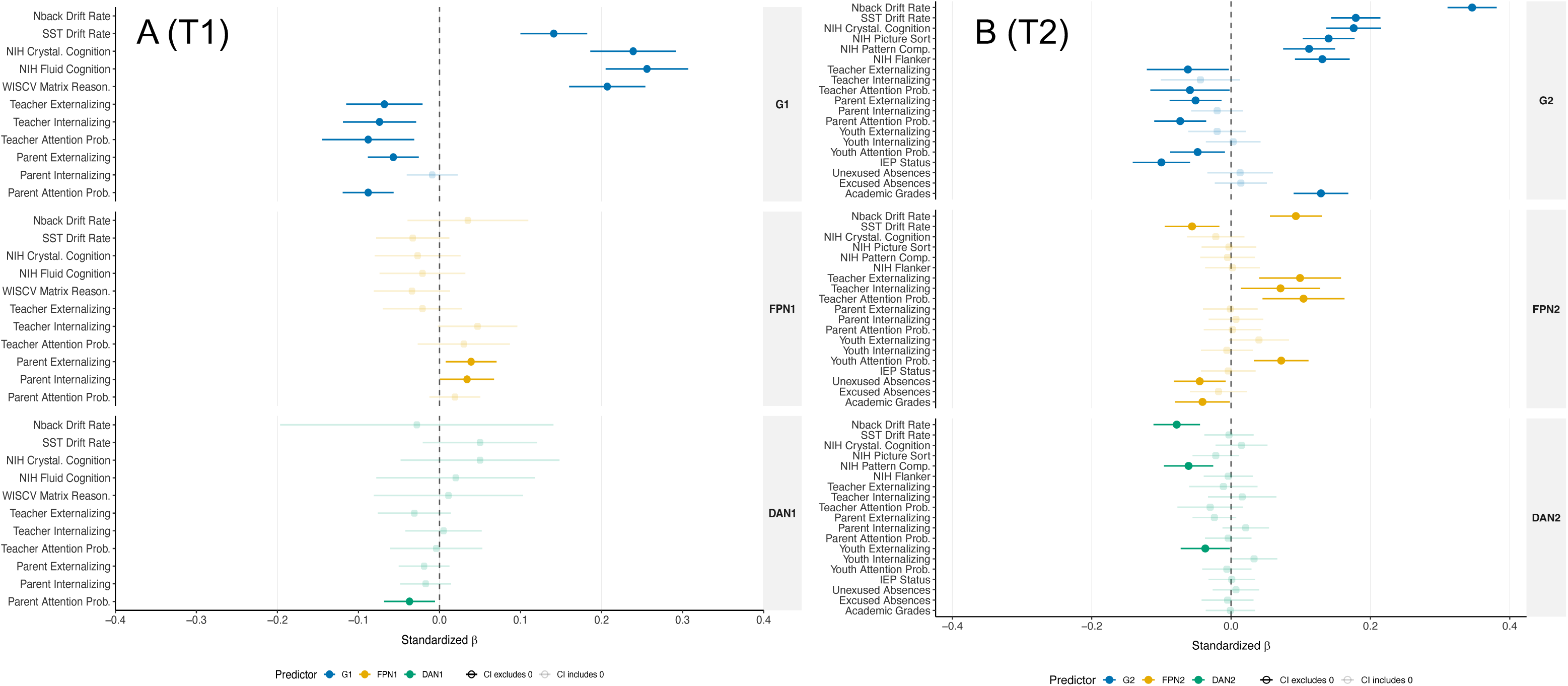
*Forest plot depicting criterion validity associations between general task-control network activation, network specific activation, and criterion measures in ABCD.* Criterion validity associations for baseline assessment depicted in panel A and criterion validity associations for 2-year follow-up depicted in panel B. T1/1 denotes baseline. T2/2 denotes 2-year follow-up. G=general task-control network activation latent factor. FPN=specific frontoparietal network latent factor. DAN=specific dorsal attention network latent factor. SST=Stop signal task. NIH=NIH toolbox cognition battery measure. Reason.=Reasoning. Crystal.=Crystallized. Prob.=Problems.

Additionally, we leveraged the ABCD dataset to test associations with broadband psychopathology symptoms, including attention problems, and school/academic functioning. We used multiple informant reports across teachers, parents, and youth where data were available.

Here again, all associations between the general factor and criterion variables were in the expected direction across both baseline and 2-year follow-up (*see* **Figure 4**). Specifically, the general factor demonstrated negative associations with all psychopathology variables across all three informant reports, such that higher general task-control network activation was related to lower scores on externalizing, internalizing, and attention problem scales. Effect sizes were all small in magnitude (β range=-0.01 to -0.09), with the largest associations observed for attention problems and the smallest for internalizing. With respect to school functioning, high general task-control network activation was associated with low likelihood of youth having an Individualize Education Program (IEP; β=-0.10) and higher academic grades (β=0.13) at 2-year follow-up. Associations between the specific FPN and DAN factors with psychopathology and school/academic functioning criterion metrics were largely not significant and small in magnitude.

In summary, results suggest that the general task-control network activation factor demonstrated a degree of stability across a two-year assessment interval in ABCD, especially after imposing strict measurement invariance. Further, this general factor demonstrated strong convergent validity across samples in the form of large, positive correlations with a variety of in- and out-of-scanner cognitive tasks, and strong criterion validity through negative correlations with psychopathology and positive correlations with academic functioning.

### Trait-state-occasion modeling reveals robust, trait-like neural signature of task-control activation

Before a claim can be made that general factors of task-control network functioning reflect trait-like, individual difference dimensions, their trait-like properties must be further examined. We harnessed the two repeated measures from ABCD to estimate a latent trait-state-occasion model of the bifactor models of task-control network functioning. This is a novel way to distinguish stable individual differences in task-control network activation from time-specific fluctuations in neural activity and measurement error. By separating these sources of variance, the model provides a more precise characterization of the stability and developmental relevance of the neural signature and clarifies whether its associations with cognitive and adaptive outcomes reflect enduring individual differences or occasion-specific variation. To our knowledge, this is the first application of a latent trait-state-occasion model to characterize the stability and occasion-specific variability of a neuroimaging feature.

The latent trait-state-occasion model provided an acceptable fit to the data (*χ*^2^(756)=10934.872, *p*<.001, CFI=.905, RMSEA=.045(0.044, 0.046), SRMR=.067). Constraining the residual variances for the general and specific occasion factors across baseline and 2-year follow-up did not lead to a significant decrement in model fit (ΔCFI=-0.001; *χ*^2^(759)=10975.931, *p*<.001, CFI=.904, RMSEA=.045(0.044, 0.046), SRMR=.064). Significant trait and occasion components emerged for general task-control network functioning and the specific FPN and DAN factors (*p*s for all factor variances <.001). Regarding general task-control network functioning, 23% of the total variance was accounted for by the trait component and 77% was accounted for by the occasion component. Similarly, trait and occasion components accounted for 25% and 75% of the variance in specific FPN activation and 27% and 73% of the variance in specific DAN activation, respectively.

Next, we examined measurement accuracy. Given the strong criterion validity demonstrated by the general factors of task-control network activation, in comparison to the specific FPN and DAN factors, in the single assessment models, we focus here on the general factor (although trait-state-occasion models of the FPN and DAN specific factors are reported in **Supplemental Material 2**). All models included head motion, and baseline age, sex, and socioeconomic status (coi_z) as covariates. The trait of general task-control network functioning had remarkably strong relationships with in- and out-of-scanner cognition, and small-to-medium relationships with psychopathology (**Figure 5**).

**Figure 5.**
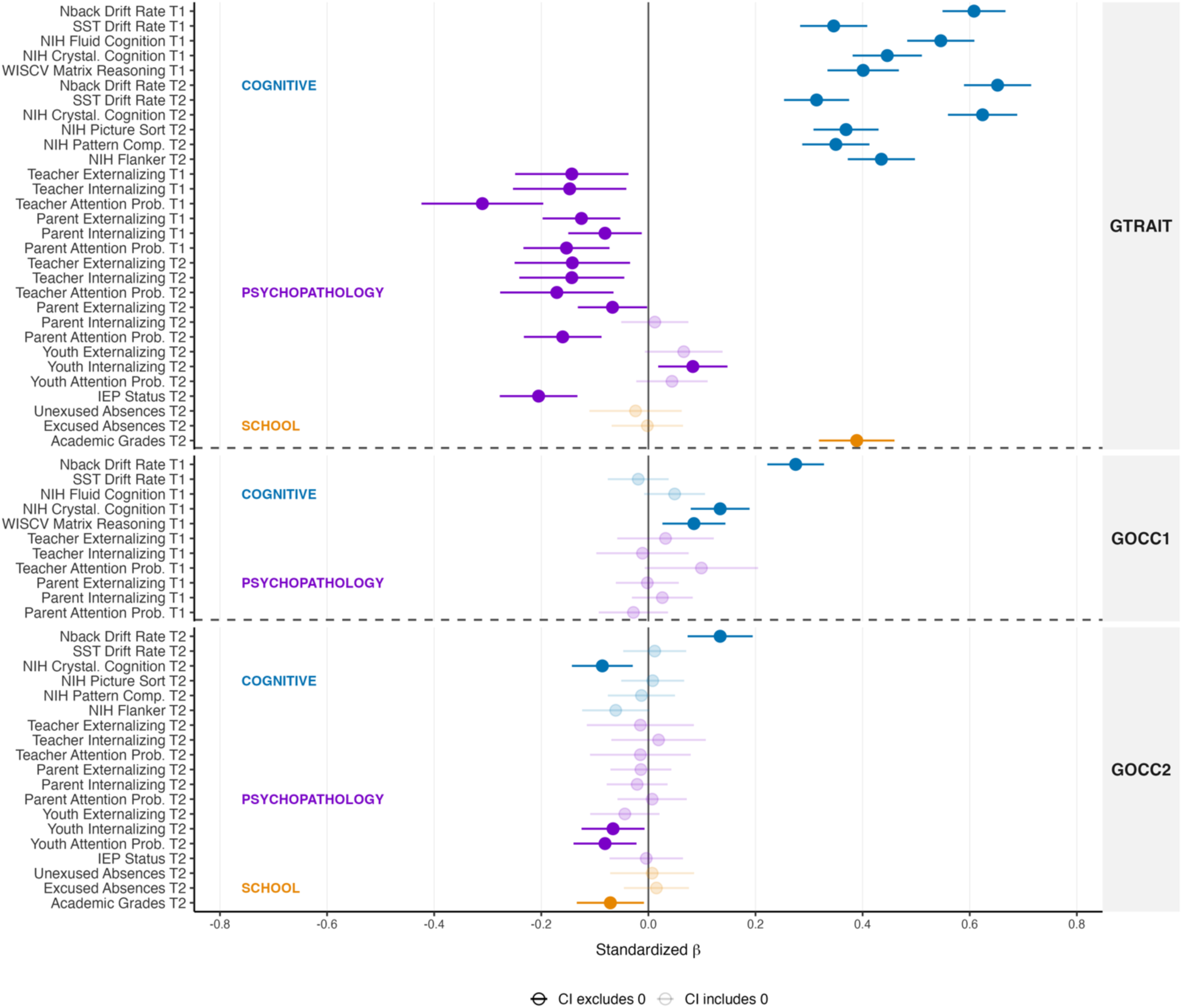
*Forest plot depicting criterion validity associations between trait and occasion factors of general task-control network activation and criterion measures*. GTRAIT=Trait of general task-control network activation latent factor. GOCC1=Baseline occasion general task-control network activation latent factor. GOCC2=2-year follow-up occasion general task-control network activation latent factor. T1=Baseline. T2=2-year follow-up. SST=Stop signal task. NIH=NIH toolbox cognition battery measure. Crystal.=Crystallized. Prob.=Problems.

Specifically, the trait general factor demonstrated strong convergent validity in the form of consistent, positive associations with n-back drift rate (β_T1_=0.61; β_T2_=0.65) and a variety of other cognitive performance measures (β range=0.31 to 0.62) that were all large in magnitude. With respect to psychopathology, the trait general factor demonstrated consistent, negative associations with broadband externalizing (β range=-0.07 to -0.14) and attention problems (β range=-0.15 to -0.31) across parent and teacher reports. Associations were less consistent with internalizing symptoms; though, most effects were small and in the negative direction. Further, we observed a large, positive association with academic grades (β=0.39) and a medium, negative association with youth having an IEP (β=-0.21). In summary, higher levels of trait general task-control network functioning were consistently related to better cognitive, academic, and school functioning and lower externalizing symptoms and clinical attention problems. These effect sizes are notably larger than most cross-modality (e.g., task behavior to questionnaire) effects that have been reported in the ABCD sample^49^ and are comparable to effect sizes reported for brain-behavior associations drawn from less parsimonious multivariate predictive models of ABCD resting state fMRI data^21^.

Occasion factors’ associations with criterion measures were generally not significant and significant associations were small in magnitude. The baseline general task-control occasion factor demonstrated a medium association with baseline concurrent drift rate (β=0.28), possibly reflecting transient fluctuations in task-control neural functioning that are associated with concurrent fluctuations in behavioral indices of neurocognition. Another possibility is that these occasion factors capture temporary context-dependent influences on neurocognition, such as sleep quality, fatigue, or arousal. To explore this further, we examined associations between the trait and occasion factors of neural activation with a range of sleep and arousal variables across both assessments. Surprisingly, associations were not significant (*see* **Supplementary Material 2; Supplementary Figure S3**), with the exception that higher general *trait* task-control neural activation was related to slightly longer parent-reported sleep duration at baseline (β=0.07) and 2-year follow-up (β=0.09).

Finally, the magnitude of associations between the trait general factor of task-control activation and cognitive, psychopathology, and school criterion metrics were notably larger in comparison to those observed with the single assessments of general task-control activation (*see* **Figure 6**). This pattern of findings indicates, for the first time, that the use of only two repeated assessments of neuroimaging can capture a neural signature of task-control network functioning that demonstrates *strong* criterion validity, likely through effectively isolating stable, accurate variance from time-specific measurement error.

**Figure 6.**
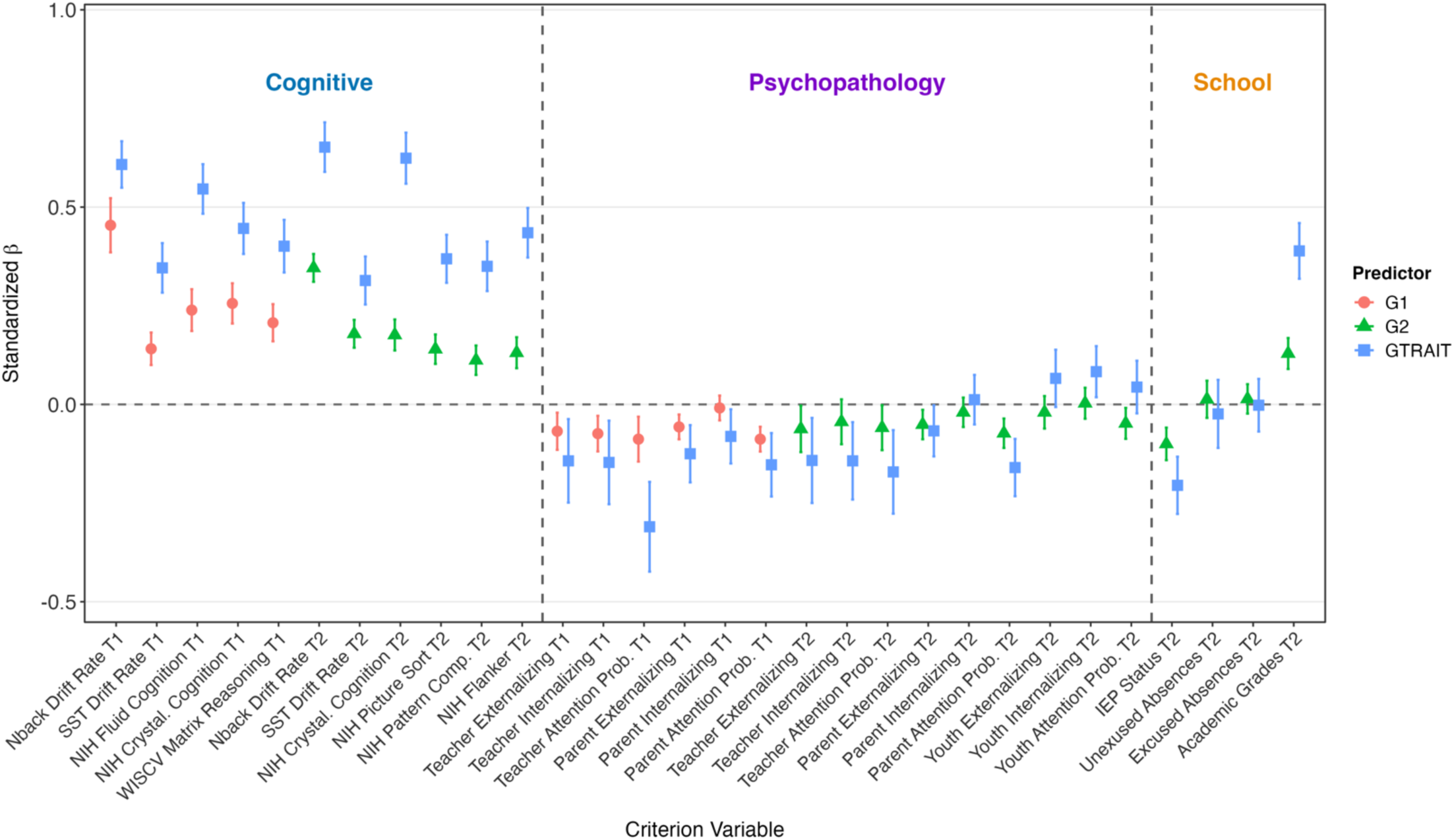
*Grouped coefficient plot comparing standardized beta coefficients for trait and state latent factors of general task-control network activation across cognitive, psychopathology, and school criterion measures.* G1=ABCD baseline state latent factor of general task-control network activation. G2=ABCD 2-year follow-up state latent factor of general task-control network activation. GTRAIT=trait latent factor of general task-control network activation. Error bars represent 95% confidence intervals. T1=Baseline. T2=2-year follow-up. SST=Stop signal task. NIH=NIH toolbox cognition battery measure. Crystal.=Crystallized. Prob.=Problems.

## Discussion

This study applied latent variable modeling to three assessments across two large neuroimaging datasets, spanning ages 9 to 37 years, to demonstrate that neural activation during n-back load in the “task-control” FPN and DAN networks is described well by a bifactor structure. Neural activation was represented by a general task-control factor that captured variance shared across the FPN and DAN, alongside two orthogonal network-specific factors.

Our findings provide strong evidence for the psychometric robustness of the general factor of task-control network activation, demonstrating that it is valid, stable, and consistently structured (i.e., invariant) across ages, samples, and scanning contexts. These findings have both substantive implications for the study of individual and developmental differences in cognitive functioning and broader methodological implications for the use of neuroimaging to characterize dimensional traits.

First, we establish a neural signature of task-control network functioning that has a stable structure from middle childhood through young adulthood. The factor structure was strictly invariant across middle childhood and early adolescence within ABCD and largely consistent when extended across independent ABCD and HCP samples spanning middle childhood to young adulthood, despite differences in participant populations, task designs, and imaging parameters. This developmentally consistent structure converges with a broader scientific effort to characterize the structure of neurocognitive abilities across development. Indeed, behavioral research has demonstrated that individual differences across a wide range of neurocognitive tasks are largely explained by domain-general latent processes across assessments and independent datasets^50–52^. Other work demonstrates longitudinal measurement invariance of a general factor of neurocognitive abilities that has been measured with both cognitive task performance and questionnaires^5,53–55^. Our findings similarly suggest that developmental changes in task-control network responses to cognitive demand may be largely quantitative, not qualitative, across critical periods of neurocognitive development. On the other hand, the increasing centrality of several prefrontal regions to the general factor in the HCP sample may point to an important qualitative developmental process that warrants further investigation. Extending our work to future ABCD data releases from young adulthood will be valuable for determining whether this pattern reflects true developmental change or sample-specific differences between ABCD and HCP.

This work represents a critical advance in developmental neuroimaging because meaningful developmental inference requires that the same latent construct be measured equivalently across age. To our knowledge, this has not yet been done in human neuroimaging. By establishing this prerequisite, our findings provide a robust framework for quantifying normative neurodevelopment, identifying atypical developmental trajectories, and evaluating how genetic and environmental factors may influence the maturation of brain features across time. Indeed, a central goal of developmental neuroscience is to understand how changes in brain structure and function give rise to age-related changes in cognition and behavior, while also elucidating how behavior and environmental experiences, in turn, shape the developing brain^56^ (e.g., dual systems theories, ^57–59^, neuroconstructivist theories^56,60^). Consequently, countless empirical studies aim to distinguish normative neurodevelopmental processes from environmentally mediated changes in neural function^61,62^, to understand how neurodevelopmental processes associate with clinical outcomes across time^63,64^, and to identify the mechanisms through which experiences influence neurocognitive outcomes^65–67^. Achieving these objectives requires neuroimaging measures that assess the same underlying constructs consistently across development, allowing observed age-related differences to be interpreted as true developmental change rather than artifacts of changing measurement properties. Although developmental neuroimaging studies routinely examine age-related differences in regional activation^16,68,69^ and connectivity^70–73^, few approaches establish that the same latent neural construct is measured equivalently across developmental stages. The neural signature established in the current study addresses this gap by providing a developmentally comparable measure of task-control network functioning, creating new opportunities to distinguish normative neurodevelopment from environmentally or clinically driven variation in neural function across time.

Second, the general task-control activation factor showed evidence of stability across a two-year measurement interval during adolescence. A prior study examined test-retest reliability of many behavioral measures of neurocognition in ABCD across the same baseline to 2-year follow-up interval^74^. Test-retest reliabilities on NIH toolbox measures ranged from fair (*r*=0.44) to excellent (*r*=0.82). Estimates were highest for composite measures, with several individual tasks displaying only fair test-retest reliabilities (*r*s<0.50). Thus, our general task-control activation factor performed comparably to established single-task behavioral measures of neurocognition. Importantly, our estimate of reliability across the two-year interval is notably higher than previous reliability estimates of individual ROIs drawn from the same task in the ABCD data set^17^, suggesting the general factor may show greater stability than the individual regions that comprise it.

Third, results from validity analyses using a wide range of criterion metrics across cognitive, psychopathology, academic/school, and sleep domains demonstrate that this general task-control activation metric is consistently linked to individuals’ cognitive abilities and an array of meaningful outcomes typically associated with cognition. Further, the effect sizes we demonstrate for associations with the trait measure are larger than most effects reported in the ABCD data set^49^ and in broader individual differences research^44^, and are comparable to those drawn from less parsimonious multivariate predictive modeling approaches applied to ABCD and HCP neuroimaging data^15,21^. Specifically, the general factor demonstrated good convergent validity in the form of large, positive associations with cognitive processing efficiency (drift rate) and medium-to-large, positive associations with a wide range of other established cognitive tasks across ABCD and HCP samples. These findings are consistent with a growing body of empirical evidence suggesting that task-control brain network responses to cognitive demand is a central neural underpinning of cognitive ability^35,36^. As prior work in this area has primarily relied on data-driven multivariate models, we advance the understanding of these network dynamics, establishing that a general factor of task-control system is consistent across development.

Consistent with prior work linking cognitive abilities to psychopathology^5,75–78^, this general factor demonstrated small-to-medium, negative associations with externalizing psychopathology and small-to-large negative correlations with attention problems across all three informant reports. Higher general task-control neural activation was also associated with better academic grades and lower likelihood of current IEP status, consistent with extensive evidence linking neurocognitive abilities to school functioning^65,79^. Further, small, positive associations between general task-control neural activation and longer sleep duration are consistent with a recent meta-analysis which found sleep loss–related impairments across a variety of neurocognitive behavioral tasks^80^, and with a meta-analysis documenting associations between sleep deprivation and decreased FPN activation^81^.

Fourth, findings from trait-state-occasion modeling revealed critical insights about the utility of repeated measurement in neuroimaging research. Specifically, the magnitude of associations between our metric of general task-control neural activation and criterion variables improved markedly when we created a trait measure spanning two time points. In contrast, the occasion factors, despite accounting for the majority of the variance in neural measures, were generally not related to criterion or state-based measures of sleep and arousal. One interpretation of these results is that the occasion factors represent construct-irrelevant measurement error. A central challenge in neuroimaging research is that observed neural measures contain substantial measurement error arising from scanner-related noise, head motion, and physiological noise (e.g., fluctuations from heart rate, respiration, blood flow, etc.)^17,82–85^. Thus, the magnitude of associations with clinical phenotypes and adaptive behavior was likely improved through effectively isolating stable, construct-relevant variance in the trait measure of general task-control neural activation from time-specific measurement error. Notably, our findings suggest that only two neuroimaging sessions may be required to statistically account for a large proportion of measurement error.

The current study also has several important methodological implications. The dominant approaches in human neuroimaging that aim to characterize the neural underpinnings of cognition rely on high-dimensional predictive models^20–23^. These models contain thousands of parameters that lack interpretability and lead to overfitting, their measurement properties and developmental stability are unknown, and, perhaps most importantly, they often fail to replicate in independent samples^21,22,29,30^. Our approach is fundamentally different, leveraging established findings from network neuroscience to guide our selection of a smaller number of ROIs and our estimation of a parsimonious, reproducible factor model. Our finding that individual differences in responses to cognitive demand across “task-positive” regions are largely driven by a single general factor is consistent with recent findings that individual differences in activations from task fMRI contrasts are typically explained by low-dimensional factor structures^16,86,87^.

Therefore, although activations across different ROIs have typically been assumed to index dissociable neural processes, factor analysis offers a critical tool for reducing high-dimensional task contrast data to more parsimonious and meaningful underlying dimensions. The current study utilized task fMRI contrast data, but as connectivity estimates from resting state fMRI also tend to show a relatively low-dimensional structure^88^, they may similarly benefit from factor analytic methods that leverage theory about largescale brain networks^89^ to characterize individual differences.

The method used in the current study yields models that are far more parsimonious and interpretable than multivariate predictive models, which typically have thousands of parameters^31–33^. Our relatively simple model showed effect sizes with criterion measures comparable to those of more complex prediction models^21^, while identifying a mechanistically interpretable dimension linked to theories of brain functioning. Whereas neural signatures from multivariate predictive models are typically criterion-defined, this signature was instead determined by the covariance structure of neural data and informed by prior findings on the functional organization of the brain. Therefore, its robust associations with criterion measures can be interpreted as relations between a substantive neural-level dimension and behavioral outcomes, rather than as simply the pattern of variance in neural-level data that shows associations with these outcomes. Finally, although multivariate predictive models are prone to overfitting data in specific samples and fail to generalize to new samples^29,30^, our relatively simpler model showed a striking evidence of generalizability, supported by formal tests of parameter equivalence across multiple age groups and two different samples with different demographic compositions, scanning protocols, and task designs. This demonstrates that theory-driven psychometric modeling can facilitate reproducibility and translatability across samples and contexts, an area of critical and growing importance to modern neuroscience^90,91^.

Several limitations should be noted. First, the ABCD and HCP samples differed in their sampling strategies, n-back task designs, and imaging parameters, which precludes strong inferences about measurement invariance and developmental differences between adolescence and young adulthood. Future applications of this approach to additional ABCD time points can strengthen inferences about developmental change. Second, generalizability of our findings is limited by the demographic composition of both samples. The geographically restricted HCP sample has limited racial and ethnic diversity, and neuroimaging quality-control procedures in ABCD may yield analytic samples that disproportionately represent White and socioeconomically advantaged youth^92,93^.

In conclusion, we demonstrate that task-control network responses to cognitive demand are largely described by a unitary factor and that developmental change in this factor across middle childhood to early adulthood is largely quantitative, not qualitative. Further, trait-state-occasion modeling using two repeated assessments in ABCD reveals a general task-control neural trait displays robust associations with a wide variety of cognitive measures, broadband psychopathology, sleep, and academic functioning, demonstrating that repeated neuroimaging assessments substantially improve validity. This neural signature provides a foundation for characterizing developmental trajectories and individual differences in goal-directed cognitive ability across the lifespan. More broadly, these findings reveal the value of combining well-validated psychometric methods with prior knowledge about brain organization to create parsimonious models of high-dimensional neuroimaging data that optimize the measurement of meaningful neural-level dimensions^16,34,95^ and offer an interpretable and reproducible alternative to the data-driven models that have become the major focus of neuroimaging research.

## Methods

### ABCD sample

The Adolescent Brain Cognitive Development^sm^ (ABCD) Study is a large, multisite study with multimodal data obtained from over 11,000 adolescents collected from 21 sites across the United States (Casey et al., 2018; Garavan et al., 2018). The study conforms to the rules and procedures of each site’s Institutional Review Board. The University of California San Diego’s IRB serves as a single IRB for the ABCD consortium (IRB#: 160091), and the IRBs of each of the 21 sites also approve the study. All participants provide informed consent (parents) or informed assent (children). High resolution structural images (1.0mm^3^ isotropic voxels) T1-weighted and T2-weighted), diffusion weighted imaging, and multiple functional MRI runs (TR=800 ms, TE=30 ms, 2.4 mm isotropic voxels) were collected for all participants.

The data included in this study came from ABCD Release 5.1 (http://dx.doi.org/10.15154/z563-zd24). The ABCD data repository grows and changes over time. Here, we included data from baseline (*M*age=10.01 years; range=8.92 to 11.00) and 2-year follow-up. (*M*age=11.99; range=10.58 to 13.75). Following exclusion criteria for neuroimaging data quality and n-back behavioral data (outlined below) the total sample at baseline was *n*=4862 and *n*=3648 were included at 2-year follow-up. Demographic statistics are summarized in **Supplementary Table S4.**

For this study, task-control network function was defined using the ABCD emotional N-back (EN-Back) task. The task is comprised of 2 different load conditions (2-back and 0-back) and the stimuli were places and happy, fearful, and neutral faces. The full task was across 2 runs of 8 blocks (four 2-back, four 0-back) each. Full details about the task design and acquisition can be found in Casey et al. (2018).

### HCP sample

The Human Connectome Project is a large dataset of *N*=1,200 healthy young adults^96^. Study procedures were approved by the Washington University Institutional Review Board (IRB#: 201204036), and all participants provided informed consent. All HCP participants included were scanned across 2 days using a customized Siemens Skyra 3T MRI scanner at Washington University in St. Louis. High resolution structural images (0.7mm^3^ isotropic voxels) T1-weighted and T2-weighted), diffusion weighted imaging, and functional MRI runs (TR=720 ms, TE=33.1 ms, 2.0 mm isotropic voxels) were collected for all participants. Following exclusion criteria outlined below, the final sample included *n*=1080 participants (*M*age=28.78 years; SD=3.69; range=22 to 37). Demographic statistics are summarized in **Supplementary Table S4.**

For this study, task-control network functioning was defined using the HCP working memory task—an N-back (2-back versus 0-back) embedded in a Category Specific (faces, places, tools, and body parts) Representation Task. Two runs (each 5 min 1 second) of this task were collected on the first day of fMRI data collection. Each block was 25 seconds, with 8 blocks (four 0-back, four 2-back) per run. Full details about the task design, acquisition, and preprocessing in HCP can be found in Barch et al. (2013)^97^. Note that the ABCD EN-back task was designed as a variant of the HCP n-back task.

### Measures

For both the ABCD EN-back and the HCP N-back tasks, the contrast of interest was 2BK-0BK to define the cortical activation of high cognitive demand.

The behavioral measures of interest in ABCD included domains across neurocognition, psychopathology, and state-based measures (sleep and arousal). The neurocognitive battery measures included the matrix reasoning task from the Wechsler Intelligence Scale for Children Fifth Edition^98^ at baseline (WISCV), and the NIH Toolbox Cognitive battery^99^ measures of fluid intelligence at baseline and crystallized intelligence at baseline and 2-year follow-up. Note that several neurocognitive composite scores were not available at 2-year follow-up due to administrative difficulties during the COVID-19 pandemic. Because of this, for the second timepoint, we instead included subscales that comprise the fluid composite score—picture sorting, pattern comparison/processing speed, and conflict processing/flanker task. More details about the tasks and composite scores used in ABCD can be found in Luciana et al., (2018)^100^. In addition to the neurocognitive battery across the WISCV and NIH Toolbox, we also included drift rate from the n-back and Stop Signal Tasks^37^, respectively. More details on these measures can be found in the *Diffusion Decision Modeling* section of the Data Analytic Strategy, below.

Psychopathology measures were continuous measures from the ASEBA scales (Achenbach & Rescorla, 2004)^101^, including parent-rated (CBCL: Child Behavior Checklist) and teacher-rated (BPM: Brief Problem Monitor)^102^ measures of externalizing, internalizing, and attention problems from baseline and 2-year follow-up. Youth provided self-reports of externalizing, internalizing, and attention problems on the Brief Problem Monitor (BPM)^102^ at 2-year follow-up.

Academic and school functioning measures were parent-reported on the School Attendance and Grades questionnaire^103^ at 2-year follow-up. Variables included current Individualized Education Plan (IEP) status (0=no; 1=yes), excused and unexcused absences in the past four weeks (0=0 days to 6=11+ days), and academic grades (reverse coded such that higher scores indicate better academic grades).

Several state-based measures were included to assess construct validity of occasion task-control network function factors. Sleep duration was assessed at baseline and 2-year follow-up using a parent-reported item from the Sleep Disturbances Scale for Children^104,105^, which was converted to an approximate hours of sleep per night. ABCD Study measures of arousal included self-reported feelings of sleepiness and tiredness assessed immediately before and after the neuroimaging protocol at baseline and 2-year follow-up^37^. On each occasion, participants were presented with a list of words describing different feelings, including “sleepy” and “tired”, and were asked to rate the extent to which they experienced each feeling in the present moment. Responses were recorded on a Likert scale ranging from 1=“Very slightly or not at all” to 5=”Extremely”.

HCP cognitive criterion measures included in-scanner drift rate, the NIH Toolbox Cognitive Battery^97,99^ measures of fluid and crystallized intelligence, and performance on the Penn Progressive Matrices ^106^ (fluid intelligence), Penn Word Memory Test^107,108^ (verbal episodic memory), and Variable Short Penn Line Orientation Test^107,108^ (spatial orientation).

In ABCD, covariates included mean framewise displacement (FD) at baseline and 2-year follow-up, and baseline age, sex, and z-scored Child Opportunity Index^109^ to account for socioeconomic status^110^. In HCP, covariates included mean FD, age, sex, and education level.

### Neuroimaging preprocessing and ROI extraction

Neuroimaging exclusion criteria, data quality checks, preprocessing, and analyses are described in detail in Weigard et al., 2024^36^. Briefly, preprocessing was performed using fMRIPrep v1.5.0^111^. Structural T1-weighted (T1w) and T2-weighted images were run through recon-all using FreeSurfer v6.0.1. Functional images were then corrected for fieldmap distortions, rigidly coregistered to the T1, motion corrected, normalized to standard space, and transformed to CIFTI space.

Task data were analyzed following HCP scripts for both datasets. Models were constructed using FSL, averaging across runs, smoothed (2mm FWHM Gaussian), and highpass filtered. First-level models included nuisance parameters for motion, white matter, cerebrospinal fluid, and framewise displacement. Each condition (2-back vs. 0-back) and category of stimuli (e.g., places) were modeled separately, then averaged across stimuli for the final relevant 2back-0back contrast. These task activation maps were then averaged within each region in the 100 parcel Schaeffer parcellation to reduce the dimensionality for factor analysis. All FPN and DAN regions from this parcellation (**Figure 1**) were retained for analysis and activation data were averaged withing each parcel. We averaged several parcels located in midline areas of the brain across hemispheres to create three bilateral variables: AVG_Cont_pCun_1 (averaging 7Networks_RH_Cont_pCun_1 and 7Networks_LH_Cont_pCun_1), AVG_Cont_Cing_1 (averaging 7Networks_RH_Cont_Cing_1 and 7Networks_LH_Cont_Cing_1), and AVG_DorsAttn_Post_5 (averaging 7Networks_RH_DorsAttn_Post_5, 7Networks_LH_DorsAttn_Post_5; *see* **Supplementary Table S3**). These corresponding left and right hemisphere parcels exhibited strong correlations with one another, likely explained by intrinsic smoothing during data collection that causes signal to bleed into adjacent areas. A total of 21 variables were included across the task-control network, 11 from the FPN and 10 from the DAN. Further details on the variables used in the current study and the corresponding naming scheme from the Schaefer 100-parcel, 7-network cortical parcellation are provided in **Supplementary Table S3.**

### Data Analytic Strategy

All Mplus model output documents and results are available on Open Science Framework at: https://osf.io/f94sw/overview?view_only=ebe2108da72e450eb934839838235fbf

#### Diffusion decision modeling

The diffusion decision model (DDM)^112,113^ was applied to the n-back task to obtain estimates of individuals’ efficiency of evidence accumulation (EEA) on the task. We used a recent version of the DDM designed to parametrically separate EEA for goal-relevant information from individuals’ strategic use of goal-irrelevant familiarity evidence to complete the task, which is important for describing individual differences in performance on the ABCD task^45^. Parameter estimation procedures were identical to those that were implemented, and described in detail, in this prior study^45^. Briefly, all models were estimated in the Dynamic Models of Choice R suite^114^ using individual-level Bayesian estimation via differential-evolution Markov chain Monte Carlo sampling^115^. Prior to estimation, trials response times less than 200ms were excluded as fast guesses and individuals with accuracy rates <0.55 or omission (i.e., non-response) rates >0.25 on either the 0-or 2-back task were excluded. Hierarchical Bayesian versions of the n-back model were first fit to a randomly selected subsample of 200 singleton ABCD participants to generate informative priors for the baseline and 2-year follow-up waves, and these individuals were then excluded from all further analyses. Individual-level Bayesian estimation using the resulting informative priors was then conducted for the rest of the ABCD sample separately at baseline and 2-year follow-up waves. Convergence was determined by confirming the Gelman-Rubin statistic of all parameters was less than 1.10^116^ and model fit was assessed using posterior predictive plots. Medians of individuals’ posterior distributions of the EEA parameter were then retained as point estimates for analyses. All procedures, initial prior distributions, and posterior predictive plots are detailed in the open access prior article and its supplement^45^.

To assess generalizability of the neural measures’ associations with EEA, we also included estimates of EEA drawn from an application of the standard DDM to baseline data from “go” trials of the ABCD stop-signal task (SST)^78^. The general procedures used for estimating this model were similar to those used for the n-back (individual-level Bayesian estimation in Dynamic Models of Choice) with the exception that broad, uninformative priors were used (*see* **Supplementary Table S5**). All main parameters of the diffusion model (*a*, *v*, *z*, *t_0_*) were estimated along with across-trial variability in in non-decision time (*s_t0_*) and a parameter representing the probability of go-failure (*P*_gf_), which captures omissions, or failures to respond due to attentional lapse^117^. Response times less than 200ms were excluded as fast guesses and individuals with accuracy rates <0.55 or omission rates >0.50 were excluded from analyses. Posterior medians of the DDM’s drift rate (*v*) parameter were retained as measures of EEA on the SST.

#### Factor analysis

To examine the bifactor structure of task-control neural activations across development, we conducted Confirmatory Factor Analysis (CFA) estimated in Mplus 8.8 using Maximum likelihood with robust standard errors (MLR)^118^. This approach allowed us to examine the factor structure and evaluate its equivalence across development and samples. A bifactor structure of task-control network activation was specified at each assessment, with all indicators loading onto a general factor, 11 indicators additionally loading onto an FPN-specific factor, and 10 additionally loading onto a DAN-specific factor (**Figure 2**). The residual variances of each indicator (ROI) were allowed to covary across both assessment points of ABCD. Residual correlations were considered when modification indices indicated substantial localized misfit. Correlations were introduced sequentially and retained when they improved model fit. This approach results in freeing 38, 33, and 27 residual correlations for the bifactor models at the ABCD baseline, ABCD 2-year follow-up, and HCP baseline assessments, respectively. Given that nested models tests are sensitive to detect small trivial differences across models when using large sample sizes, we used change in model comparative fit index (CFI) > .01 to indicate significant change in model fit^119,120^. Fit of the CFAs was assessed using conventional (e.g., CFI, root mean square error approximation (RMSEA), standardized root mean square residual (SRMR)) as well as alternative fit statistics developed for bifactor models (e.g., omega, omega hierarchical subscale (ωHS), and construct replicability (H))^121–123^. ω and ωs are the latent variable analogues to coefficient alpha for the general task-positive network factor and specific FPN and DAN factors. ωH differs from ω in that it only represents the variance from a single latent variable whereas ω is a function of all common variance. ωHS is an index of the reliability of the specific factors after partitioning out the variance attributable to the general factor. The only computational difference between ωS and ωHS is that in the numerator, the term associated with variance on the general factor is removed, leaving only the variance associated with the specific factor. ω, ωS, ωH, and ωHS values greater than .70 indicate acceptable reliability and values greater than .80 indicate good reliability^122,124–126^. To evaluate fit of the bifactor model, values lower than .80 for Percentage of Uncontaminated Variance (PUC) and greater than .70 for Explained Common Variance (ECV) indicate that data are unidimensional enough to introduce bias in fitting a multidimensional model^43,127^. Values of construct replicability, H, >.70 indicate that measured items adequately represent the latent factors^123^.

Next, we used single assessment bifactor models to examine associations with a wide array of criterion measures. Criterion measures were grouped by construct (e.g., cognitive ability, psychopathology, academic/school functioning) and reporter (parent, teacher, youth), with each category estimated in separate models to reduce model complexity. Within each model, criterion measures were allowed to covary (e.g., teacher-reported externalizing, internalizing, and attention problems). In ABCD, youth were clustered within families and stratified by study site using the CLUSTER IS and STRATIFICATION IS statements, respectively. In HCP, individuals were clustered within families using the CLUSTER IS statement.

#### Latent trait-state-occasion modeling

Strong evidence of measurement invariance allowed us to leverage the two repeated measures from ABCD to conduct latent trait-state-occasion modeling to examine the trait-like properties of the task-control network functioning factors. As described by Cole and colleagues (2005)^128^, the model was specified such that the latent variables at each of the two time points (i.e., the state factors) loaded onto a time-invariant (trait) factor, with the factor loadings fixed to one. In addition, time-variant (occasion) factors were created that were defined by the corresponding latent variable from that time point, with factor loadings fixed to one (e.g., Time-variant general task-control factor at Time 1 was defined by the general task-control factor variable at Time 1). The residual variances of the state factors were fixed to zero to distribute all variance to either time-invariant or time-variant factors. Factor loadings were held to equality for each indicator across assessments points, consistent with the previously demonstrated longitudinal invariance. Given that we only included two assessments, autoregressive time-variant paths were not included to facilitate model identification. To account for shared method variance across time, the residual variances of each indicator were allowed to covary across assessment points, as in the longitudinal measurement models described previously^129^. Factor means were set to zero. This structure was specified for the general task-control network factors as well as the specific FPN and DAN factors, such that each construct had trait, state, and occasion factors.

## Acknowledgements

Data used in the preparation of this article were obtained from the Adolescent Brain Cognitive Development (ABCD) Study (https://abcdstudy.org), held in the NIMH Data Archive (NDA). This is a multisite, longitudinal study designed to recruit more than 10,000 children aged 9-10 years and follow them over 10 years into early adulthood. The ABCD Study is supported by the National Institutes of Health and additional federal partners under award numbers U01DA041022, U01DA041028, U01DA041048, U01DA041089, U01DA041106, U01DA041117, U01DA041120, U01DA041134, U01DA041148, U01DA041156, U01DA041174, U24DA041123, and U24DA041147. A full list of supporters is available at https://abcdstudy.org/nih-collaborators. A listing of participating sites and a complete listing of the study investigators can be found at https://abcdstudy.org/principal-investigators.html. ABCD consortium investigators designed and implemented the study and/or provided data but did not necessarily participate in analysis or writing of this report. This manuscript reflects the views of the authors and may not reflect the opinions or views of the NIH or ABCD consortium investigators. The ABCD data repository grows and changes over time. The ABCD data used in this report came from NDA Study 576, DOI 10.15154/1412097, which can be found at https://ndar.nih.gov/study.html?id=576. Additional data were provided by the Human Connectome Project, WU-Minn Consortium (Principal Investigators: David Van Essen and Kamil Ugurbil; 1U54MH091657) funded by the 16 NIH Institutes and Centers that support the NIH Blueprint for Neuroscience Research; and by the McDonnell Center for Systems Neuroscience at Washington University. We thank Cleanthis Michael for his assistance in generating an icon of the frontoparietal and dorsal attention networks used in the graphical abstract.

## References

1. Strenze, T. Intelligence and socioeconomic success: A meta-analytic review of longitudinal research. Intelligence 35, 401–426 (2007).

2. Caspi, A., Moffitt, T. & Poulton, R. Lifelong Impact of Early Self-Control. Am. Sci. 101, 352 (2013).

3. Moffitt, T. E., et al. A gradient of childhood self-control predicts health, wealth, and public safety. Proc. Natl. Acad. Sci. U.S.A. 108, 2693–2698 (2011).

4. Hofer, S. M. & Clouston, S. On the Importance of Early-Life Cognitive Abilities in Shaping Later-Life Outcomes. Research in Human Development 11, 241–246 (2014).

5. Paige, K. J., Meisel, S. N. & Colder, C. R. An examination of reciprocal associations between substance use and effortful control across adolescence using a bifactor model of externalizing symptoms. Dev Psychopathol 33, 1507–1519 (2021).

6. Stucke, N. J. & Doebel, S. Early childhood executive function predicts concurrent and later social and behavioral outcomes: A review and meta-analysis. Psychological Bulletin 150, 1178–1206 (2024).

7. Deer, L. K., Hastings, P. D. & Hostinar, C. E. The Role of Childhood Executive Function in Explaining Income Disparities in Long-Term Academic Achievement. Child Development 91, e1046–e1063 (2020).

8. Mill, R. D., Ito, T. & Cole, M. W. From connectome to cognition: The search for mechanism in human functional brain networks. NeuroImage 160, 124–139 (2017).

9. Kanai, R. & Rees, G. The structural basis of inter-individual differences in human behaviour and cognition. Nature Reviews Neuroscience 12, 231–242 (2011).

10. Braver, T. S., Cole, M. W. & Yarkoni, T. Vive les differences! Individual variation in neural mechanisms of executive control. Current opinion in neurobiology 20, 242–250 (2010).

11. Crone, E. A. & Steinbeis, N. Neural perspectives on cognitive control development during childhood and adolescence. Trends in cognitive sciences 21, 205–215 (2017).

12. Luna, B., Padmanabhan, A. & O’Hearn, K. What has fMRI told us about the development of cognitive control through adolescence? Brain and cognition 72, 101–113 (2010).

13. Volkow, N. D. et al. The conception of the ABCD study: From substance use to a broad NIH collaboration. Developmental cognitive neuroscience 32, 4–7 (2018).

14. Luna, B., Padmanabhan, A. & O’Hearn, K. What has fMRI told us about the Development of Cognitive Control through Adolescence? Brain and Cognition 72, 101–113 (2010).

15. Vieira, B. H. et al. On the prediction of human intelligence from neuroimaging: A systematic review of methods and reporting. Intelligence 93, 101654 (2022).

16. Paige, K. J. et al. Clarifying the longitudinal factor structure, temporal stability, and construct validity of Go/No-Go task-related neural activation across adolescence and young adulthood. Developmental Cognitive Neuroscience 67, 101390 (2024).

17. Kennedy, J. T. et al. Reliability and stability challenges in ABCD task fMRI data. NeuroImage 252, 119046 (2022).

18. Elliott, M. L. et al. What Is the Test-Retest Reliability of Common Task-Functional MRI Measures? New Empirical Evidence and a Meta-Analysis. Psychol Sci 31, 792–806 (2020).

19. Dubois, J. & Adolphs, R. Building a science of individual differences from fMRI. Trends in cognitive sciences 20, 425–443 (2016).

20. Shearer, C. B. A resting state functional connectivity analysis of human intelligence: Broad theoretical and practical implications for multiple intelligences theory. Psychology & Neuroscience 13, 127–148 (2020).

21. Marek, S. et al. Reproducible brain-wide association studies require thousands of individuals. Nature 603, 654–660 (2022).

22. Dubois, J., Galdi, P., Paul, L. K. & Adolphs, R. A distributed brain network predicts general intelligence from resting-state human neuroimaging data. Phil. Trans. R. Soc. B 373, 20170284 (2018).

23. Rosenberg, M. D. et al. A neuromarker of sustained attention from whole-brain functional connectivity. Nature neuroscience 19, 165–171 (2016).

24. Zdorovtsova, N. et al. The entropy of resting-state neural dynamics is a marker of general cognitive ability in childhood. Preprint at 10.1101/2023.08.08.552448 (2023).

25. Sripada, C. et al. Prediction of neurocognition in youth from resting state fMRI. Mol Psychiatry 25, 3413–3421 (2020).

26. Fox, M. D., Corbetta, M., Snyder, A. Z., Vincent, J. L. & Raichle, M. E. Spontaneous neuronal activity distinguishes human dorsal and ventral attention systems. Proc. Natl. Acad. Sci. U.S.A. 103, 10046–10051 (2006).

27. Mulder, M. J., Van Maanen, L. & Forstmann, B. U. Perceptual decision neurosciences – A model-based review. Neuroscience 277, 872–884 (2014).

28. Machner, B. et al. Resting-State Functional Connectivity in the Dorsal Attention Network Relates to Behavioral Performance in Spatial Attention Tasks and May Show Task-Related Adaptation. Front. Hum. Neurosci. 15, 757128 (2022).

29. Lanka, P. et al. Supervised machine learning for diagnostic classification from large-scale neuroimaging datasets. Brain imaging and behavior 14, 2378–2416 (2020).

30. Janssen, R. J., Mourão-Miranda, J. & Schnack, H. G. Making individual prognoses in psychiatry using neuroimaging and machine learning. Biological psychiatry: Cognitive neuroscience and neuroimaging 3, 798–808 (2018).

31. Yarkoni, T. & Westfall, J. Choosing prediction over explanation in psychology: Lessons from machine learning. Perspectives on Psychological Science 12, 1100–1122 (2017).

32. Lipton, Z. C. The mythos of model interpretability. Communications of the ACM 61, 36– 43 (2018).

33. Kohoutová, L. et al. Toward a unified framework for interpreting machine-learning models in neuroimaging. Nature protocols 15, 1399–1435 (2020).

34. Cooper, S. R., Jackson, J. J., Barch, D. M. & Braver, T. S. Neuroimaging of individual differences: A latent variable modeling perspective. Neuroscience & Biobehavioral Reviews 98, 29–46 (2019).

35. Sripada, C., Angstadt, M., Rutherford, S., Taxali, A. & Shedden, K. Toward a “treadmill test” for cognition: Improved prediction of general cognitive ability from the task activated brain. Human Brain Mapping 41, 3186–3197 (2020).

36. Weigard, A. et al. Flexible adaptation of task-positive brain networks predicts efficiency of evidence accumulation. Commun Biol 7, 801 (2024).

37. Casey, B. J. et al. The Adolescent Brain Cognitive Development (ABCD) study: Imaging acquisition across 21 sites. Developmental Cognitive Neuroscience 32, 43–54 (2018).

38. Van Essen, D. C. et al. The WU-Minn Human Connectome Project: An overview. NeuroImage 80, 62–79 (2013).

39. Vincent, J. L., Kahn, I., Snyder, A. Z., Raichle, M. E. & Buckner, R. L. Evidence for a frontoparietal control system revealed by intrinsic functional connectivity. Journal of neurophysiology (2008).

40. Avelar-Pereira, B., Bäckman, L., Wåhlin, A., Nyberg, L. & Salami, A. Age-related differences in dynamic interactions among default mode, frontoparietal control, and dorsal attention networks during resting-state and interference resolution. Frontiers in Aging Neuroscience 9, 152 (2017).

41. Spreng, R. N., Sepulcre, J., Turner, G. R., Stevens, W. D. & Schacter, D. L. Intrinsic architecture underlying the relations among the default, dorsal attention, and frontoparietal control networks of the human brain. Journal of cognitive neuroscience 25, 74–86 (2013).

42. Godbersen, G. M. et al. Task-evoked metabolic demands of the posteromedial default mode network are shaped by dorsal attention and frontoparietal control networks. Elife 12, e84683 (2023).

43. Reise, S. P. The Rediscovery of Bifactor Measurement Models. Multivariate Behavioral Research 47, 667–696 (2012).

44. Gignac, G. E. & Szodorai, E. T. Effect size guidelines for individual differences researchers. Personality and Individual Differences 102, 74–78 (2016).

45. Weigard, A. et al. Dissociable Contributions of Goal-Relevant Evidence and Goal-Irrelevant Familiarity to Individual and Developmental Differences in Conflict Recognition. Cognitive Science 48, e70019 (2024).

46. Tomlinson, R. C. et al. Efficiency of evidence accumulation as a formal model-based measure of task-general executive functioning in adolescents. Journal of psychopathology and clinical science (2025).

47. Weigard, A., Clark, D. A. & Sripada, C. Cognitive efficiency beats top-down control as a reliable individual difference dimension relevant to self-control. Cognition 215, 104818 (2021).

48. Marek, S. et al. Patterns of brain-wide associations reflect socioeconomics. Science 392, eaee6213 (2026).

49. Owens, M. M. et al. Recalibrating expectations about effect size: A multi-method survey of effect sizes in the ABCD study. PloS one 16, e0257535 (2021).

50. Tervo-Clemmens, B. et al. A canonical trajectory of executive function maturation from adolescence to adulthood. Nat Commun 14, 6922 (2023).

51. Löffler, C., Frischkorn, G. T., Hagemann, D., Sadus, K. & Schubert, A.-L. The common factor of executive functions measures nothing but speed of information uptake. Psychological Research 88, 1092–1114 (2024).

52. Vermeent, S., Young, E. S., DeJoseph, M. L., Schubert, A. & Frankenhuis, W. E. Cognitive deficits and enhancements in youth from adverse conditions: An integrative assessment using Drift Diffusion Modeling in the ABCD study. Developmental Science 27, (2024).

53. Willoughby, M. T., Wirth, R. J., Blair, C. B., & Family Life Project Investigators. Executive function in early childhood: Longitudinal measurement invariance and developmental change. Psychological Assessment 24, 418–431 (2012).

54. Reilly, S. E., Downer, J. T. & Grimm, K. J. Developmental trajectories of executive functions from preschool to kindergarten. Developmental Science 25, e13236 (2022).

55. Castellanos-Ryan, N. et al. Modelling executive function across early childhood: Longitudinal invariance, development from 3.5 to 7 years and later academic performance. Cognitive Development 68, 101365 (2023).

56. Johnson, M. H. & De Haan, M. Developmental Cognitive Neuroscience: An Introduction. (Wiley Blackwell, Chichester, West Sussex, 2015).

57. Meisel, S. N., Fosco, W. D., Hawk, L. W. & Colder, C. R. Mind the gap: A review and recommendations for statistically evaluating Dual Systems models of adolescent risk behavior. Developmental Cognitive Neuroscience 39, 100681 (2019).

58. Shulman, E. P. et al. The dual systems model: Review, reappraisal, and reaffirmation. Developmental Cognitive Neuroscience 17, 103–117 (2016).

59. Steinberg, L. A dual systems model of adolescent risk-taking. Developmental Psychobiology 52, 216–224 (2010).

60. Karmiloff-Smith, A. Nativism versus neuroconstructivism: Rethinking the study of developmental disorders. Developmental Psychology 45, 56–63 (2009).

61. Kim-Spoon, J. et al. Maltreatment and brain development: The effects of abuse and neglect on longitudinal trajectories of neural activation during risk processing and cognitive control. Developmental Cognitive Neuroscience 48, 100939 (2021).

62. Sheridan, M. A. & McLaughlin, K. A. Introduction to the special issue on childhood adversity and neurodevelopment. Developmental Cognitive Neuroscience 54, 101082 (2022).

63. Romer, A. L. & Pizzagalli, D. A. Is executive dysfunction a risk marker or consequence of psychopathology? A test of executive function as a prospective predictor and outcome of general psychopathology in the adolescent brain cognitive development study®. Developmental Cognitive Neuroscience 51, 100994 (2021).

64. Jensen, C. D., Jacola, L. M. & Schatz, J. Introduction to the Special Issue on Neural Processes and Pediatric Health: Creating a Pediatric Health Neuroscience Research Agenda. Journal of Pediatric Psychology 43, 815–820 (2018).

65. Paige, K. J. et al. Leveraging bifactor modeling to test prospective direct and indirect effects of adolescent alcohol use and externalizing symptoms on the development of task-general executive functioning. Dev Psychopathol 1–22 (2024) doi:10.1017/S095457942400138X.

66. Hong, S.-J. et al. Decomposing complex links between the childhood environment and brain structure in school-aged youth. Developmental Cognitive Neuroscience 48, 100919 (2021).

67. Zhang, Y. L. et al. Childhood Material Hardship Linked to Adolescent Neurocognition: A Computational Modeling Approach. Developmental Science 29, e70213 (2026).

68. Luna, B., Garver, K. E., Urban, T. A., Lazar, N. A. & Sweeney, J. A. Maturation of Cognitive Processes From Late Childhood to Adulthood. Child Development 75, 1357–1372 (2004).

69. Ordaz, S. J., Foran, W., Velanova, K. & Luna, B. Longitudinal Growth Curves of Brain Function Underlying Inhibitory Control through Adolescence. J. Neurosci. 33, 18109–18124 (2013).

70. Vinette, S. A. & Bray, S. Variation in functional connectivity along anterior-to-posterior intraparietal sulcus, and relationship with age across late childhood and adolescence. Developmental Cognitive Neuroscience 13, 32–42 (2015).

71. Marek, S., Hwang, K., Foran, W., Hallquist, M. N. & Luna, B. The Contribution of Network Organization and Integration to the Development of Cognitive Control. PLoS Biol 13, e1002328 (2015).

72. Barber, A. D., Caffo, B. S., Pekar, J. J. & Mostofsky, S. H. Developmental changes in within- and between-network connectivity between late childhood and adulthood. Neuropsychologia 51, 156–167 (2013).

73. Kardan, O. et al. Differences in the functional brain architecture of sustained attention and working memory in youth and adults. PLoS Biol 20, e3001938 (2022).

74. Anokhin, A. P. et al. Age-related changes and longitudinal stability of individual differences in ABCD Neurocognition measures. Developmental Cognitive Neuroscience 54, 101078 (2022).

75. Brislin, S. J. et al. Differentiated nomological networks of internalizing, externalizing, and the general factor of psychopathology (‘p factor’) in emerging adolescence in the ABCD study. Psychological medicine 52, 3051–3061 (2022).

76. Blanken, L. M. E. et al. Cognitive functioning in children with internalising, externalising and dysregulation problems: a population-based study. Eur Child Adolesc Psychiatry 26, 445– 456 (2017).

77. Blok, E. et al. Cognitive performance in children and adolescents with psychopathology traits: A cross-sectional multicohort study in the general population. Dev Psychopathol 35, 926–940 (2023).

78. Weigard, A. et al. Generalizable prediction of childhood ADHD symptoms from neurocognitive testing and youth characteristics. Transl Psychiatry 13, 225 (2023).

79. Valiente, C., Lemery-Chalfant, K., Swanson, J. & Reiser, M. Prediction of children’s academic competence from their effortful control, relationships, and classroom participation. Journal of Educational Psychology 100, 67–77 (2008).

80. Cao, Y., Xie, T. & Ma, N. The impairments of sleep loss on core executive functions: General and task-specific effects. Sleep Medicine Reviews 84, 102163 (2025).

81. Ma, N., Dinges, D. F., Basner, M. & Rao, H. How Acute Total Sleep Loss Affects the Attending Brain: A Meta-Analysis of Neuroimaging Studies. Sleep 38, 233–240 (2015).

82. Greve, D. N. et al. A Survey of the Sources of Noise in fMRI. Psychometrika 78, 396–416 (2013).

83. Elliott, M. L. et al. What Is the Test-Retest Reliability of Common Task-Functional MRI Measures? New Empirical Evidence and a Meta-Analysis. Psychol Sci 31, 792–806 (2020).

84. Hoeppli, M. E. et al. Denoising task-related FMRI : Balancing noise reduction against signal loss. Human Brain Mapping 44, 5523–5546 (2023).

85. Liu, T. T. Noise contributions to the fMRI signal: An overview. NeuroImage 143, 141–151 (2016).

86. Cooper, S. R., Jackson, J. J., Barch, D. M. & Braver, T. S. Neuroimaging of individual differences: A latent variable modeling perspective. Neuroscience & Biobehavioral Reviews 98, 29–46 (2019).

87. Demidenko, M. I., Mumford, J. A., Ram, N. & Poldrack, R. A. A multi-sample evaluation of the measurement structure and function of the modified monetary incentive delay task in adolescents. Developmental Cognitive Neuroscience 65, 101337 (2024).

88. Sripada, C. et al. Basic units of inter-individual variation in resting state connectomes. Scientific reports 9, 1900 (2019).

89. Gates, K. M., Fisher, Z. F. & Bollen, K. A. Latent variable GIMME using model implied instrumental variables (MIIVs). Psychological methods 25, 227 (2020).

90. Falk, E. B. et al. What is a representative brain? Neuroscience meets population science. Proceedings of the National Academy of Sciences 110, 17615–17622 (2013).

91. Yarkoni, T. The generalizability crisis. Behavioral and Brain Sciences 45, e1 (2022).

92. Gard, A. M., Hyde, L. W., Heeringa, S. G., West, B. T. & Mitchell, C. Why weight? Analytic approaches for large-scale population neuroscience data. Developmental Cognitive Neuroscience 59, 101196 (2023).

93. Cosgrove, K. T. et al. Limits to the generalizability of resting-state functional magnetic resonance imaging studies of youth: An examination of ABCD Study® baseline data. Brain Imaging and Behavior 16, 1919–1925 (2022).

94. Constantinidis, C. & Luna, B. Neural Substrates of Inhibitory Control Maturation in Adolescence. Trends in Neurosciences 42, 604–616 (2019).

95. Moriarity, D. P. & Alloy, L. B. Back to Basics: The Importance of Measurement Properties in Biological Psychiatry. Neuroscience & Biobehavioral Reviews 123, 72–82 (2021).

96. Van Essen, D. C. et al. The WU-Minn Human Connectome Project: An overview. NeuroImage 80, 62–79 (2013).

97. Barch, D. M. et al. Function in the human connectome: Task-fMRI and individual differences in behavior. NeuroImage 80, 169–189 (2013).

98. Wechsler, D. Wechsler Intelligence Scale for Children--Fifth Edition. 10.1037/t79359-000 (2024).

99. Weintraub, S. et al. Cognition assessment using the NIH Toolbox. Neurology 80, (2013).

100. Luciana, M. et al. Adolescent neurocognitive development and impacts of substance use: Overview of the adolescent brain cognitive development (ABCD) baseline neurocognition battery. Developmental Cognitive Neuroscience 32, 67–79 (2018).

101. Achenbach, T. M., & Rescorla, L. A. Manual for the ASEBA School-Age Forms & Profiles. (Burlington, VT: University of Vermont, Research Center for Children, Youth, & Families., 2001).

102. Thomas M. Achenbach. Achenbach System of Empirically Based Assessment (ASEBA): Development, Findings, Theory, and Applications. (University of Vermont, Research Center of Children, Youth & Families.).

103. Zucker, R. A. et al. Assessment of culture and environment in the Adolescent Brain and Cognitive Development Study: Rationale, description of measures, and early data. Developmental Cognitive Neuroscience 32, 107–120 (2018).

104. Bruni, O. et al. The Sleep Disturbance Scale for Children (SDSC) Construct ion and validation of an instrument to evaluate sleep disturbances in childhood and adolescence. Journal of Sleep Research 5, 251–261 (1996).

105. Ferreira, V. R. et al. Sleep Disturbance Scale for Children: Translation, cultural adaptation, and validation. Sleep Medicine 10, 457–463 (2009).

106. Bilker, W. B. et al. Development of Abbreviated Nine-Item Forms of the Raven’s Standard Progressive Matrices Test. Assessment 19, 354–369 (2012).

107. Gur, R. C. et al. A cognitive neuroscience-based computerized battery for efficient measurement of individual differences: Standardization and initial construct validation. Journal of Neuroscience Methods 187, 254–262 (2010).

108. Gur, R. Computerized Neurocognitive Scanning: I. Methodology and Validation in Healthy People. Neuropsychopharmacology 25, 766–776 (2001).

109. Fan, C. C. et al. Adolescent Brain Cognitive Development (ABCD) study Linked External Data (LED): Protocol and practices for geocoding and assignment of environmental data. Developmental Cognitive Neuroscience 52, 101030 (2021).

110. Marek, S. et al. Patterns of brain-wide associations reflect socioeconomics. Science 392, eaee6213 (2026).

111. Esteban, O. et al. fMRIPrep: a robust preprocessing pipeline for functional MRI. Nat Methods 16, 111–116 (2019).

112. Ratcliff, R. A theory of memory retrieval. Psychological Review 85, 59–108 (1978).

113. Ratcliff, R., Smith, P. L., Brown, S. D. & McKoon, G. Diffusion Decision Model: Current Issues and History. Trends in Cognitive Sciences 20, 260–281 (2016).

114. Heathcote, A. et al. Dynamic models of choice. Behav Res 51, 961–985 (2019).

115. Turner, B. M., Sederberg, P. B., Brown, S. D. & Steyvers, M. A method for efficiently sampling from distributions with correlated dimensions. Psychological Methods 18, 368–384 (2013).

116. Gelman, A. & Rubin, D. B. Inference from Iterative Simulation Using Multiple Sequences. Statist. Sci. 7, (1992).

117. Damaso, K. A. M. et al. A cognitive model of response omissions in distraction paradigms. Mem Cogn 50, 962–978 (2022).

118. Muthén, L.K., & Muthén, B.O. Mplus User’s Guide. Eighth Edition. (Muthén & Muthén., Los Angeles, CA, 1998).

119. Cheung, G. W. & Rensvold, R. B. Evaluating Goodness-of-Fit Indexes for Testing Measurement Invariance. Structural Equation Modeling: A Multidisciplinary Journal 9, 233– 255 (2002).

120. Paige, K. J. et al. Leveraging bifactor modeling to test prospective direct and indirect effects of adolescent alcohol use and externalizing symptoms on the development of task-general executive functioning. Dev Psychopathol 1–22 (2024) doi:10.1017/S095457942400138X.

121. Dueber, D. M. Bifactor Indices Calculator. 473 KB University of Kentucky Libraries 10.13023/EDP.TOOL.01 (2017).

122. Rodriguez, A., Reise, S. P. & Haviland, M. G. Applying Bifactor Statistical Indices in the Evaluation of Psychological Measures. J Pers Assess 98, 223–237 (2016).

123. Rodriguez, A., Reise, S. P. & Haviland, M. G. Evaluating bifactor models: Calculating and interpreting statistical indices. Psychol Methods 21, 137–150 (2016).

124. Cortina, J. M. What is coefficient alpha? An examination of theory and applications. Journal of Applied Psychology 78, 98–104 (1993).

125. Santos, J. R. A. Cronbach’s Alpha: A Tool for Assessing the Reliability of Scales. Journal of Extension 37, (1999).

126. Tavakol, M. & Dennick, R. Making sense of Cronbach’s alpha. Int. J. Medical Education 2, 53–55 (2011).

127. Quinn, H. O. Bifactor Models, Explained Common Variance (ECV), and the Usefulness of Scores from Unidimensional Item Response Theory Analyses. https://doi.org/10.17615/T6FF-A088 doi:10.17615/T6FF-A088.

128. Cole, D. A., Martin, N. C. & Steiger, J. H. Empirical and Conceptual Problems With Longitudinal Trait-State Models: Introducing a Trait-State-Occasion Model. Psychological Methods 10, 3–20 (2005).

129. LaGrange, B. & Cole, D. A. An Expansion of the Trait-State-Occasion Model: Accounting for Shared Method Variance. Structural Equation Modeling: A Multidisciplinary Journal 15, 241–271 (2008).

