## Supplemental Material 1 for "A parsimonious, age-invariant, and psychometrically validated neural signature of task-control network functioning"

*Supplementary Table S1*. Fit statistics for single assessment bifactor models.

| Model | $\chi 2$ (df), *p* value | CFI | RMSEA | SRMR |
|---|---|---|---|---|
| ABCD baseline | | | | |
| Initial Model | 16138.615(169), <.001 | 0.706 | 0.139(0.138, 0.141) | 0.083 |
| Allowing 38 residual correlations | 4595.815(131), <.001 | 0.918 | 0.084(0.082, 0.086) | 0.050 |
| ABCD 2-year follow-up | | | | |
| Initial Model | 11040.460(169), <.001 | 0.722 | 0.133(0.131, 0.135) | 0.083 |
| Allowing 33 residual correlations | 3123.083(136), <.001 | 0.924 | 0.078(0.075, 0.080) | 0.056 |
| HCP | | | | |
| Initial Model | 2695.239(169), <.001 | 0.780 | 0.118(0.114, 0.122) | 0.065 |
| Allowing 27 residual correlations | 1476.927(150), <.001 | 0.885 | 0.091(0.086, 0.095) | 0.048 |

*Note*. We freed residual correlations when modification indices indicated substantial localized misfit and when they improved model fit. Across both the ABCD 2-year follow-up and HCP models, the LH_DorsAttn_Post_4 indicator produced a negative, not significant residual variance. Thus, we set these residual variances to zero. ABCD=Adolescent Brain Cognitive Development Study. HCP=Human Connectome Project. CFI=comparative fit index. RMSEA=root mean square error approximation. SRMR=standardized root mean square residual.

*Supplementary Table S2*. Fit statistics for structural equation models examining construct validity.

| Model | $\chi2$ (df), *p* value | CFI | RMSEA | SRMR |
|---|---|---|---|---|
| ABCD baseline | | | | |
| Cognitive measures | 5395.945(301), <.001 | 0.920 | 0.059(0.057, 0.060) | 0.056 |
| Teacher-reported psychopathology | 4217.111(261), <.001 | 0.930 | 0.057(0.056, 0.059) | 0.048 |
| Parent-reported psychopathology | 4208.952(261), <.001 | 0.933 | 0.056(0.054, 0.057) | 0.046 |
| ABCD 2-year follow-up | | | | |
| Cognitive measures | 4492.801(333), <.001 | 0.908 | 0.059(0.057, 0.060) | 0.059 |
| Teacher-reported psychopathology | 3740.648(276), <.001 | 0.913 | 0.060(0.058, 0.062) | 0.054 |
| Parent-reported psychopathology | 3716.204(276), <.001 | 0.918 | 0.059(0.057, 0.060) | 0.053 |
| Youth-reported psychopathology | 3732.144(276), <.001 | 0.915 | 0.059(0.057, 0.060) | 0.053 |
| Academic and school functioning | 3849.678(295), <.001 | 0.911 | 0.058(0.056, 0.059) | 0.053 |
| HCP | | | | |
| Cognitive measures | 2314.725(336), <.001 | 0.883 | 0.070(0.067, 0.073) | 0.066 |
| State-trait-occasion model | | | | |
| Cognitive measures | 16326.808(1416), <.001 | 0.895 | 0.031(0.031, 0.032) | 0.066 |
| Teacher-reported psychopathology | 12102.170(1201), <.001 | 0.900 | 0.032(0.031, 0.032) | 0.058 |
| Parent-reported psychopathology | 12006.723(1210), <.001 | 0.919 | 0.029(0.028, 0.029) | 0.054 |
| Youth-reported psychopathology | 11567.586(1087), <.001 | 0.902 | 0.031(0.030, 0.031) | 0.057 |
| Academic and school functioning | 11891.422(1125), <.001 | 0.899 | 0.031(0.030, 0.031) | 0.058 |
| State-based sleep and arousal measures | 13012.115(1368), <.001 | 0.922 | 0.028(0.028, 0.029) | 0.052 |

*Note*. ABCD=Adolescent Brain Cognitive Development Study. HCP=Human Connectome Project. CFI=comparative fit index. RMSEA=root mean square error approximation. SRMR=standardized root mean square residual. All models included age, sex, head motion, and a measure of socioeconomic status (coi_z for ABCD and education level for HCP).

*Supplementary Table S3*. Naming scheme for regions of interest derived from Schaefer 100-parcel, 7-network cortical parcellation.

| Current Study Variable | Schaefer 100-parcel, 7-network cortical parcellation label | Network Name | Anatomical Location |
|---|---|---|---|
| RH_Cont_Par_1 | 7Networks_RH_Cont_Par_1 | control | right inferior parietal lobule |
| RH_Cont_Par_2 | 7Networks_RH_Cont_Par_2 | control | right posterior parietal lobe |
| RH_Cont_PFCl_1 | 7Networks_RH_Cont_PFCl_1 | control | right inferior lateral prefrontal cortex |
| RH_Cont_PFCl_2 | 7Networks_RH_Cont_PFCl_2 | control | |
| RH_Cont_PFCl_3 | 7Networks_RH_Cont_PFCl_3 | control | |
| RH_Cont_PFCl_4 | 7Networks_RH_Cont_PFCl_4 | control | right superior frontal gyrus |
| RH_Cont_PFCmp_1 | 7Networks_RH_Cont_PFCmp_1 | control | |
| LH_Cont_Par_1 | 7Networks_LH_Cont_Par_1 | control | left superior parietal lobule |
| LH_Cont_PFCl_1 | 7Networks_LH_Cont_PFCl_1 | control | left middle dorsolateral prefrontal cortex |
| AVG_Cont_pCun_1 | 7Networks_RH_Cont_pCun_1<br>7Networks_LH_Cont_pCun_1 | control | average of right and left precuneus |
| AVG_Cont_Cing_1 | 7Networks_RH_Cont_Cing_1<br>7Networks_LH_Cont_Cing_1 | control | average of right and left posterior cingulate cortex |
| RH_DorsAttn_Post_1 | 7Networks_RH_DorsAttn_Post_1 | dorsal attention | right lateral temporooccipital cortices |
| RH_DorsAttn_Post_3 | 7Networks_RH_DorsAttn_Post_3 | dorsal attention | right intraparietal sulcus/superior parietal lobule area |
| RH_DorsAttn_Post_4 | 7Networks_RH_DorsAttn_Post_4 | dorsal attention | right dorsal posterior parietal lobe |
| RH_DorsAttn_PrCv_1 | 7Networks_RH_DorsAttn_PrCv_1 | dorsal attention | |
| RH_DorsAttn_FEF_1 | 7Networks_RH_DorsAttn_FEF_1 | dorsal attention | right frontal eye field |
| LH_DorsAttn_Post_3 | 7Networks_LH_DorsAttn_Post_3 | dorsal attention | left intraparietal sulcus/superior parietal lobule area |
| LH_DorsAttn_Post_4 | 7Networks_LH_DorsAttn_Post_4 | dorsal attention | left postcentral sulcus |
| LH_DorsAttn_PrCv_1 | 7Networks_LH_DorsAttn_PrCv_1 | dorsal attention | left ventral precentral gyrus |
| LH_DorsAttn_FEF_1 | 7Networks_LH_DorsAttn_FEF_1 | dorsal attention | left frontal eye field |
| AVG_DorsAttn_Post_5 | 7Networks_RH_DorsAttn_Post_5<br>7Networks_LH_DorsAttn_Post_5 | dorsal attention | average of right and left posterior parietal cortex |

*Note*. Several parcels located in midline areas of the brain were also averaged across hemispheres to create three bilateral variables, denoted by "AVG". These corresponding left and right hemisphere parcels exhibited strong correlations with one another, likely explained by smoothing in the data that causes signal to bleed into adjacent areas.

*Supplementary Table S4*. Demographic details of ABCD and HCP neuroimaging samples.

| | ABCD Baseline (N = 4862) | | ABCD 2-Year (N = 3648) | | HCP (N = 1080) | |
|---|---|---|---|---|---|---|
| | N | % | N | % | N | % |
| Biological sex | | | | | | |
| Female | 2353 | 48.40 | 1832 | 50.22% | 586 | 54.26 |
| Male | 2508 | 51.58 | 1816 | 49.78% | 494 | 45.74 |
| Missing | 1 | 0 | 0 | 0 | 0 | 0 |
| Age (years) | M=10.01 | SD=0.62 | M=11.99 | SD=0.66 | M=28.78 | SD=3.69 |
| (Parent) Education: | | | | | | |
| < HS Diploma | 130 | 2.67 | 143 | 3.92 | 34 | 3.15 |
| HS Diploma/GED | 298 | 6.13 | 253 | 6.94 | 151 | 13.98 |
| Some College | 1122 | 23.08 | 897 | 24.59 | 268 | 24.81 |
| Bachelor | 1374 | 28.26 | 990 | 27.14 | 455 | 42.13 |
| Post Grad Degree | 1939 | 39.88 | 1362 | 37.34 | 170 | 15.74 |
| Missing | 4 | 0.08 | 3 | 0.08 | 2 | 0.19 |
| Race: | | | | | | |
| White | 3976 | 81.78 | 2920 | 80.04 | 812 | 75.19 |
| Black | 538 | 11.07 | 448 | 12.28 | 155 | 14.35 |
| Other | 159 | 3.27 | 141 | 3.87 | 47 | 4.35 |
| Asian | 113 | 2.32 | 78 | 2.14 | 64 | 5.93 |
| AIAN/NHPI | 30 | 0.62 | 20 | 0.55 | 2 | 0.19 |
| Missing | 46 | 0.95 | 41 | 1.12 | 0 | 0 |

*Note*. Parent education is provided for ABCD. Participant education is provided for HCP.

*Supplementary Table S5*. Broad, uninformative priors used for the parameters of the diffusion model fit to the stop-signal task data in ABCD.

| parameter | bounds | μ | σ |
|---|---|---|---|
| *a* | 0, ∞ | 1.000 | 0.500 |
| *v* | -∞, ∞ | 3.000 | 1.000 |
| *z* | 0, 1 | 0.500 | 0.100 |
| *t0* | 0, 2 | 0.300 | 0.100 |
| *st0* | 0, 2 | 0.100 | 0.050 |
| $p_{gf}$ | -∞, ∞ | 0.000 | 1.000 |

*Note*. All priors were truncated normal distributions defined by mean (μ) and scale (σ) parameters and bounded at the reported values. The $p_{gf}$ parameter is estimated on the probit scale using this prior and then converted to the natural scale as a probability.
