## Supplemental Material 2 for "A parsimonious, age-invariant, and psychometrically validated neural signature of task-control network functioning"

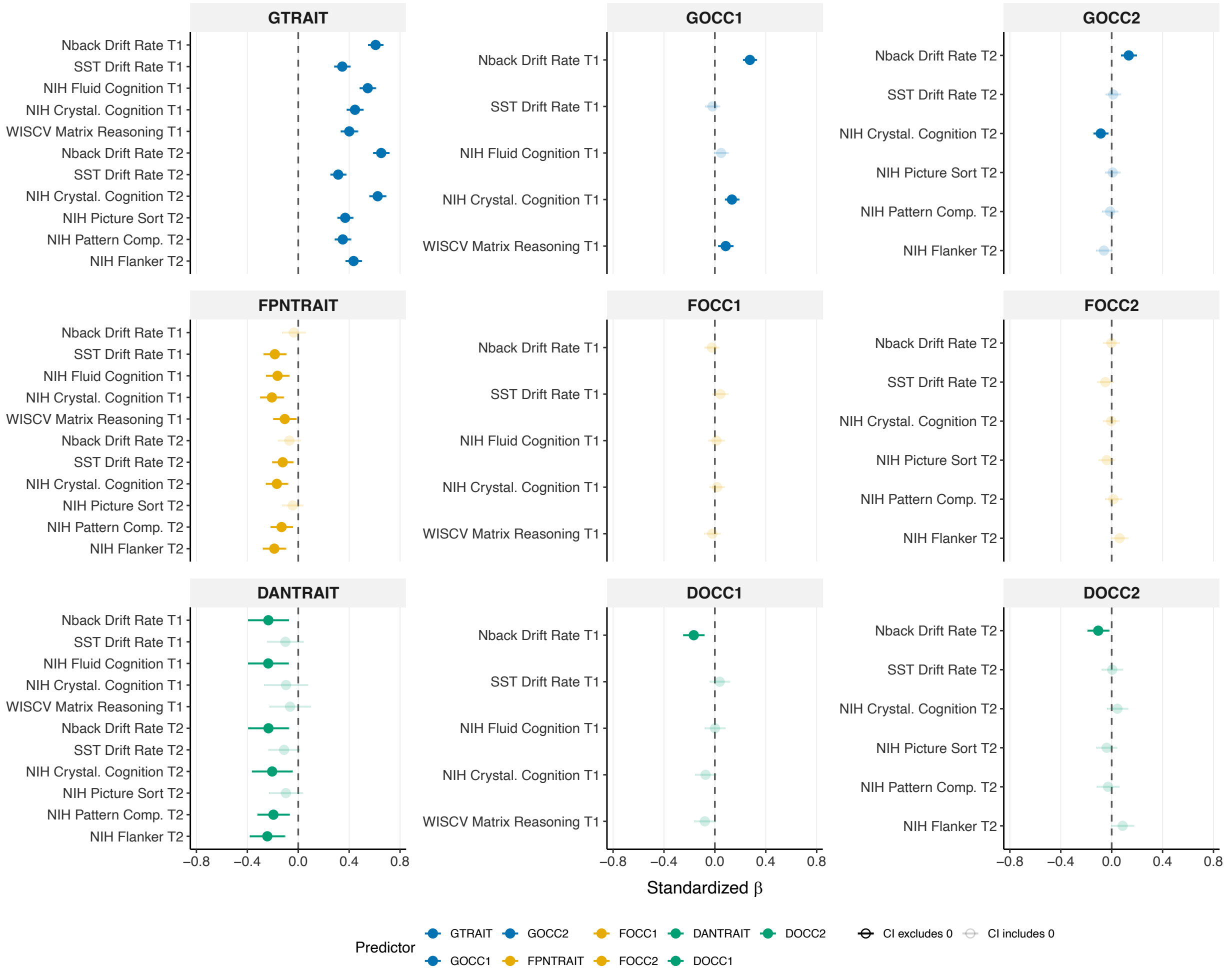


Supplementary Figure 1. Forest plot depicting the convergent validity of the trait and occasion factors for general task-control network activation, specific frontoparietal network activation, and specific dorsal attention network activation.

*Results from convergent validity analysis of network-specific trait and occasion factors*

The trait of specific FPN activation showed several small, negative associations with out-of-scanner cognitive measures. Specifically, the trait specific FPN factor was related to cognitive processing efficiency on the stop signal task ($\beta_{T1}$ = -0.18; $\beta_{T2}$ = -0.12), as well as several NIH toolbox measures across baseline (fluid composite β = -0.16; crystallized composite β = -0.21) and 2-year follow-up (crystallized composite β = -0.17; pattern comparison β = -0.13; flanker task β = -0.19). The occasion factors for specific FPN activation showed no significant associations with cognitive measures.

Similarly, the trait of specific DAN activation demonstrated small, negative relationships with in- and out-of-scanner cognitive measures. Neural activation that was unique to the DAN (nonshared with the general task-control activation) was related to lower scores on in-scanner cognitive processing efficiency ($\beta_{T1}$ = -0.24; $\beta_{T2}$ = -0.23), and several NIH toolbox measures across baseline (fluid composite β = -0.24) and 2-year follow-up (crystallized composite β = -0.21; pattern comparison β = -0.20; flanker task β = -0.24). The occasion factors showed small, negative associations with in-scanner drift rate at baseline ($\beta$ = -0.17) and 2-year follow-up ($\beta$ = -0.11).

In summary, results suggest that youth who had higher activation in FPN and DAN that was *separate* from general task-control network activation displayed *poorer* performance on behavioral cognitive tasks.

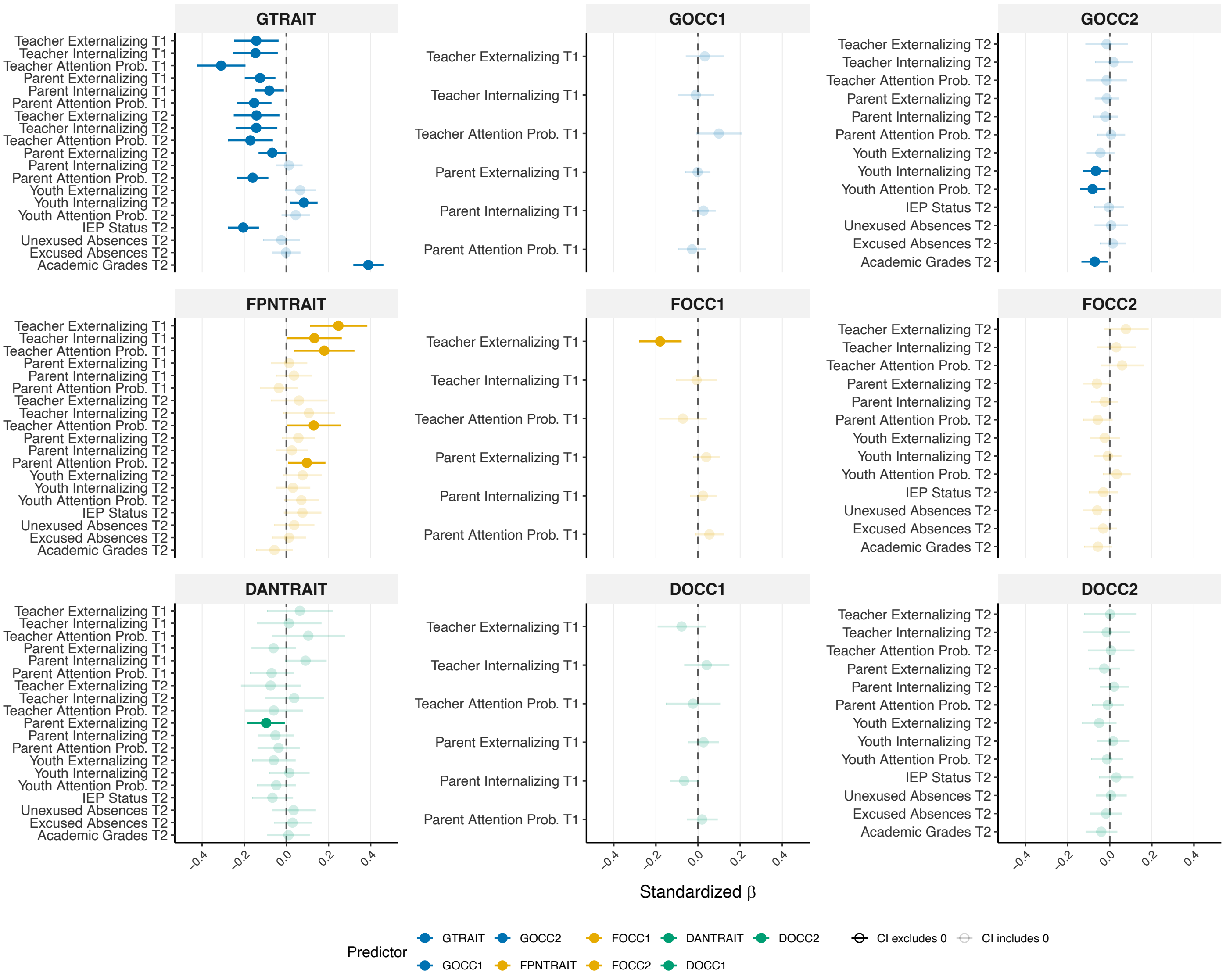


Supplementary Figure 2. Forest plot depicting the criterion validity of the trait and occasion factors for general task-control network activation, specific frontoparietal network activation, and specific dorsal attention network activation.

*Results from criterion validity analysis of network-specific trait and occasion factors*

Associations between the trait and occasion specific FPN factors and criterion measures were largely not significant. The trait of specific FPN activation showed positive associations with teacher-reported externalizing ($\beta_{T1} = 0.25$), internalizing ($\beta_{T1} = 0.13$), and attention ($\beta_{T1} = 0.18$; $\beta_{T2} = 0.13$) problems, and parent-reported attention problems ($\beta_{T2} = 0.10$). Regarding the occasion specific FPN factors, there was only one significant association. At baseline, the occasion specific FPN factor was negatively related to teacher-reported externalizing problems ($\beta = -0.18$).

The trait and occasion specific DAN factors demonstrated no significant associations with psychopathology and academic/school functioning criterion measures, with the exception of one small association between the trait specific DAN factor and parent-reported externalizing problems ($\beta$ =-0.10).

In summary, the trait factor for the specific FPN factor showed small associations with psychopathology, such that youth who had higher activation in trait FPN that was separate from general task-control network activation displayed higher levels of psychopathology symptoms. The trait and occasion factors for the specific DAN neural functioning showed largely not significant associations with psychopathology and academic/school functioning criterion measures.

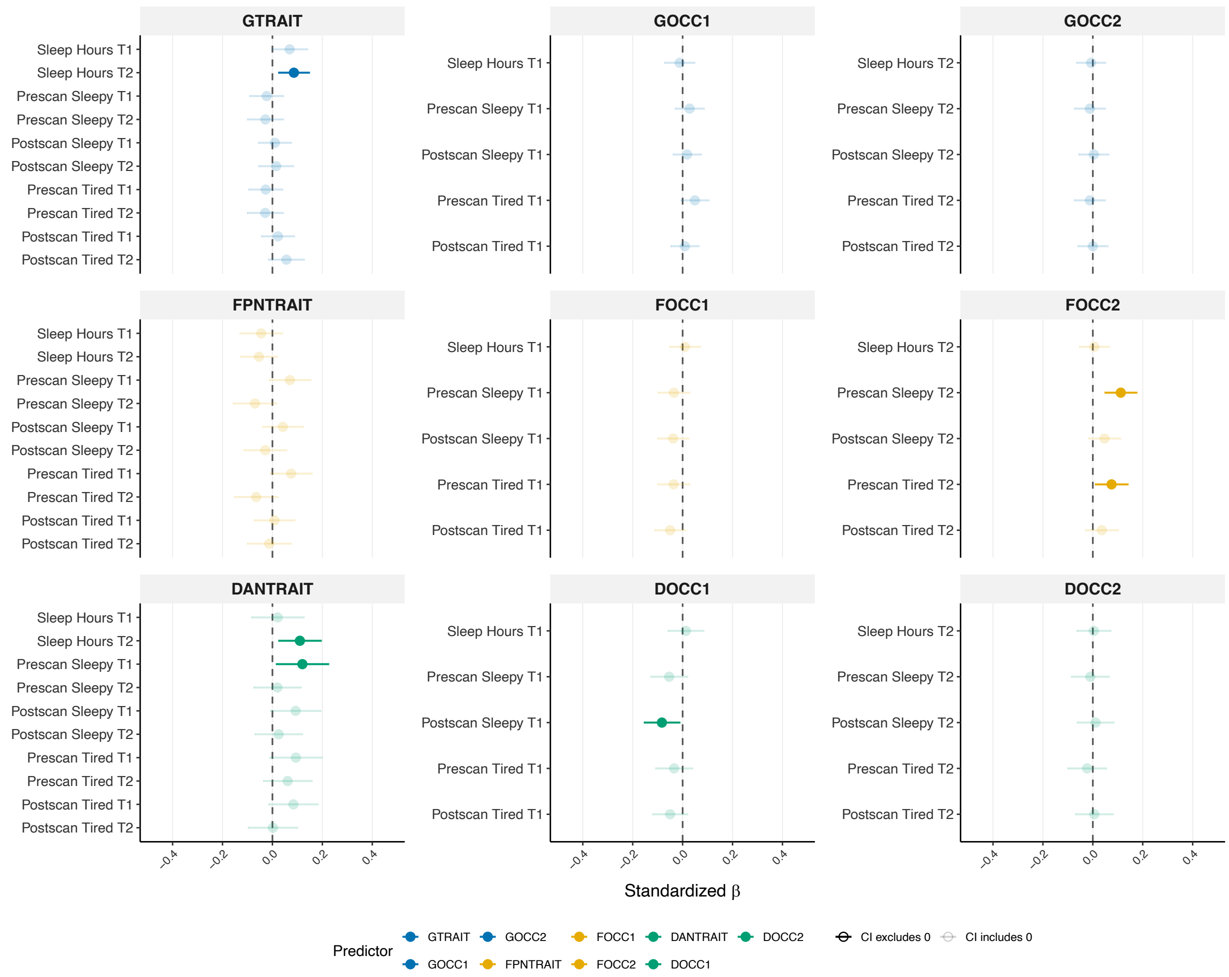


Supplementary Figure 3. Forest plot depicting associations between the trait and occasion factors for general task-control network activation, specific frontoparietal network activation, and specific dorsal attention network activation and state measures of sleep and arousal.

*Results from criterion validity analysis of task-control trait and occasion factors and state-based measures.*

Associations were largely not significant across the general and specific factors at both the state and occasion levels. The general trait factor of task-control neural activation demonstrated two small, positive associations with average hours of sleep per night (parent-reported) at baseline ($\beta = 0.07$) and 2-year follow-up ($\beta = 0.09$). No significant associations between the occasion factors for general task-control activation and state-based criterion measures were observed.

The trait measure of specific FPN neural activation demonstrated no significant associations with state-based criterion measures. The occasion measure of specific FPN neural activation at 2-year follow-up was associated with youth-rated feelings of sleepiness pre-scan ($\beta = 0.11$) scan, and feelings of tiredness pre-scan ($\beta = 0.08$). Youth who reported greater sleepiness and tiredness during this assessment demonstrated higher occasion-specific FPN activation, indicating that transient states at the time of scanning may influence neural activation patterns unique to the FPN that are not captured by the stable trait component of FPN activation. However, we observed no associations between the occasion factor for specific FPN activation and state-based criterion measures at baseline; thus, results should be interpreted cautiously.

The trait specific DAN factors demonstrated two small associations between sleep duration at 2-year follow-up ($\beta = 0.11$) and pre-scan sleepiness at baseline ($\beta = 0.12$). The occasion specific DAN factors demonstrated largely not significant associations, with the exception of one small association between the baseline occasion specific DAN factor and post-scan feelings of sleepiness ($\beta = -0.08$).
